# Establishment of plant lines cured of vertically transmitted ‘good’ viruses: The effects on plant growth and seed characteristics

**DOI:** 10.64898/2026.09.27.754801

**Authors:** Satish B. Viswanathan, Alex M. Murphy, Adrienne E. Pate, Sofia L. Miccolis, Leonas M. Engelhardt, John P. Carr

## Abstract

Several pathogenic viruses benefit plants under specific circumstances, e.g., by inducing drought resistance. It is hypothesised that RNA viruses in the *Endornaviridae* and *Partitiviridae* families, sometimes called ‘persistent’ since they are exclusively seed-transmitted at ∼100% efficiency, have followed evolutionary trajectories towards mutualism. However, evidence comes from crossing non-identical host backgrounds making it impossible to distinguish definitively between the effects of viral versus host genotypes. We used virus-induced gene silencing (VIGS) to cure pepper plants of bell pepper endornavirus (BPEV) and the partitiviruses pepper cryptic virus 1 (PCV1) and PCV2 and compared these with genetically identical plants retaining these viruses. We found that PCV1, PCV2 and BPEV can significantly affect aerial and root growth and fruit/seed characteristics in ways that could benefit host fitness. VIGS-mediated curing allows unambiguous hypothesis-testing of potential benefits of persistent viruses and exploration of how viral and host genotypes shape persistent virus–host relationships.

## INTRODUCTION

Most well-studied plant viruses are pathogenic and predominantly transmitted horizontally by vectors or wounding. Although some can be transmitted vertically through seed or pollen, most do so at low rates due to their exclusion from meristematic tissue by a combination of salicylate- and RNA silencing-mediated resistance mechanisms (Hoffmann and Incarbone, 2024; Liu and Ding, 2024; MacDiarmid et al., 2026). However, plant viruses exist that are exclusively vertically transmitted at very high rates via seed (∼100%) and pollen (30–100%) (Boccardo et al. 1987; Arancibia et al., 1995; Valverde & Gutierrez, 2008; Hershlag et al., 2020). Generally, these viruses engender no obvious disease symptoms, and it is hard to distinguish between plants of infected and uninfected lineages. Roossinck (2013) proposed the term ‘persistent’ for these inherited viruses on account of their extremely high rates of vertical transmission. It was hypothesised that these viruses persist inter-generationally because they perform useful functions for their hosts, i.e., they might be considered ‘good’ viruses (Roossinck, 2011). Particles of the first-discovered persistent viruses, the partitiviruses beet cryptic virus 1 (BCV1) and BCV2, were detected fortuitously using electron microscopy in certain sugar beet (*Beta vulgaris* var. *saccharifera*) lines (Pullen, 1968). Neither BCV1 nor BCV2 exerted any clear phenotypic effects on virus-carrying plants (Xie et al., 1994).

Plant persistent viruses occur in the RNA virus families *Amalgaviridae*, *Chrysoviridae*, *Endornaviridae*, *Partitiviridae*, and *Totiviridae* (Adams et al., 2014; Roossinck, 2010, 2013), of which the best studied are endornaviruses and partitiviruses. Endornaviruses accumulate cytoplasmically as non-enveloped double-stranded (ds) genomic RNA complexed with the viral RNA-directed RNA polymerase (RdRp). Despite accumulating as dsRNA, phylogenetic analyses indicate that endornaviruses evolved from single-stranded positive-sense RNA viruses (Gibbs et al., 2000; Valverde et al., 2019). Endornavirus RNAs have a single open reading frame encoding a presumed polyprotein. Contrastingly, partitiviruses accumulate cytoplasmically as true virus particles possessing a T=1 icosahedral shell of 60 coat protein dimers. Partitivirus genomes are bipartite: genomic RNA 1 encodes the viral RdRp, whilst RNA 2 encodes the coat protein. These genomic RNAs are packaged separately (Byrne et al., 2021).

Persistent viruses are widespread in wild plants and occur in certain lines of crops including, pepper, legumes, rice, cucurbits, and barley (Horiuchi et al., 2001; Coutts, 2005; Candresse et al., 2016; Nordenstedt et al., 2017; Khankhum et al., 2018; Mutuku et al., 2018; Okada et al., 2018). As mentioned above, persistent viruses cause no obvious disease, leading to the hypothesis that they are mutualistic symbionts (Roossinck, 2012; Fukuhara, 2019; Takahashi et al., 2020). Some evidence supports this. For example, pepper plants (*Capsicum annuum*) carrying either the partitivirus pepper cryptic virus 1 (PCV1) or bell pepper endornavirus (BPEV) are provided with some protection against aphids (Safari et al., 2019; Paudel et al., 2024). Another instance of protection from insect herbivory was provided by a study of the wild legume *Trifolium repens* (white clover). Plants carrying the partitivirus white clover cryptic virus 1 (WCCV1) were protected from fungus gnats, possibly due to virus-induced changes in volatile organic compound emission (van Molken et al., 2012). Nakatsukasa-Akune et al. (2005) found that the WCCV1 coat protein sequence is identical to that of *T. repens Early Nodulin Downregulation 1* (*TrEnodDR1*), suggestive of virus-to-host horizontal gene transfer. In white clover *TrEnodDR1* is downregulated during nodulation and expressing it constitutively in transgenic *Lotus japonicus* suppressed nodulation (Nakatsukasa-Akune et al., 2005; Suzuki et al., 2001). This suggested that during white clover’s evolution it co-opted the WCCV1 coat protein as a regulatory factor for establishing symbiosis with nitrogen-fixing bacteria (Nakatsukasa-Akune et al., 2005).

Persistent viruses may affect plant reproduction. Common bean (*Phaseolus vulgaris*) lines from the Black Turtle Soup background harbouring mixed infections of Phaseolus vulgaris endornavirus 1 (PvEV1) and PvEV2 produced longer seed pods carrying seeds of greater mass than plants of non-infected lines (Khankhum and Valverde, 2018). However, in several other common bean backgrounds the effects on seed yield or mass of PvEV1, PvEV2 or PvEV3 were unclear (Brine, Viswanathan et al., 2023). With respect to the effects of persistent viruses, studies of the amalgavirus southern tomato virus (STV) in tomato (*Solanum lycopersicum*) yielded ambiguous data. STV was suggested to increase accumulation of the pathogenic virus cucumber mosaic virus (CMV) at early timepoints and increase the severity of synergistic symptoms in mixed infections of CMV and pepino mosaic virus (González et al, 2021). However, STV also appeared to increase tomato seed production (Fukuhara et al., 2020). Escalante and Valverde (2019) hypothesised that BPEV is mildly parasitic to pepper as it appeared to inhibit plant growth, decrease seed production and diminish germination rates. In *Vicia faba*, a double-stranded RNA, later identified as Vicia faba endornavirus, engendered male sterility (Grill and Garger, 1981).

The least ambiguous results on how persistent viruses modify plant phenotypes have been obtained using lines developed by the Valverde lab (Escalante and Valverde, 2019; Khankhum and Valverde, 2018; Safari et al., 2019; Escalante et al., 2023; Paudel et al., 2024). They crossed sub-varieties of Black Turtle Soup common bean doubly infected with PvEV1 and PvEV2 with uninfected lines. They also crossed varieties of pepper with or without either PCV1 or BPEV and then performed five back-crosses (based on Escalante and Valverde, 2019) to obtain nearly isogenic infected and non-infected lines. Unfortunately, crossing has limitations in that it is time-consuming, impractical to carry out for multiple lines of plants and cannot be used to selectively remove a persistent virus from its native host plant background. Importantly, crossing cannot produce infected and non-infected plants that are completely isogenic. Even after repeated backcrossing the phenomenon of linkage drag may occur with unpredictable effects on hybrid phenotypes (Stam & Zeven, 1981); effects that might also include responses to persistent viruses. Isogenic lines of plants that either carry or are free of persistent virus infection are essential to unambiguously test if plant-persistent virus relationships are consistently mutualistic. In this work we used virus-induced gene silencing (VIGS) to establish isogenic lines of pepper plants infected with or cured of PCV1, PCV2 or BPEV.

## RESULTS

### Screening pepper cultivars for partitivirus and endornavirus infection

Using reverse transcription-coupled PCR (RT-PCR) with appropriate primers (Supplementary Table 1), twenty cultivars belonging variously to the pepper species *C. annuum*, *C. baccatum*, *C. chinense*, and *C. frutescens* (Supplementary Table 2) were screened for the presence of known pepper endornaviruses and partitiviruses (Supplementary Table 3 and Supplementary Figure 1a and b). Eight of these cultivars were infected singly with either BPEV, PCV1 or PCV2 and one line, Gourmet Jalapeno, was co-infected with PCV1 and PCV2. Sequences of the RT-PCR amplicons are shown in Supplementary Table 4 and examples of RT-PCR results shown in Supplementary Figure 1a and b. Viral RNAs were present at similar levels in all tissues tested (Supplementary Figure 1c). Full- or near-full-length genome sequences of persistent viruses from seven cultivars were obtained either by high throughput sequencing (HTS) using Illumina and metagenomic analysis (Californian Wonder, Red King, Mirasol Chilli, Gourmet Jalapeno, Poupila, and Hungarian Wax) or by RT-PCR followed by Nanopore sequencing (Cayenne Red) and deposited in GenBank (Supplementary Table 5).

Detecting BPEV in the Californian Wonder and Red King varieties was not unexpected. BPEV had been reported previously in Californian Wonder, and both lines are bell pepper cultivars, in which this endornavirus is widespread (Okada et al., 2011; Safari and Roossinck, 2018). Consistent with previous findings, HTS revealed that the BPEV sequences in Red King and Californian Wonder shared 99.68% nucleotide identity and clustered within the same major clade identified in phylogenetic analyses based on full-length and near full-length genome sequences (Tomašechová et al. 2019; Galipienso et al., 2021; Jo et al., 2022;) (Figure 1a; Supplementary Table 6).

**Figure 1.**
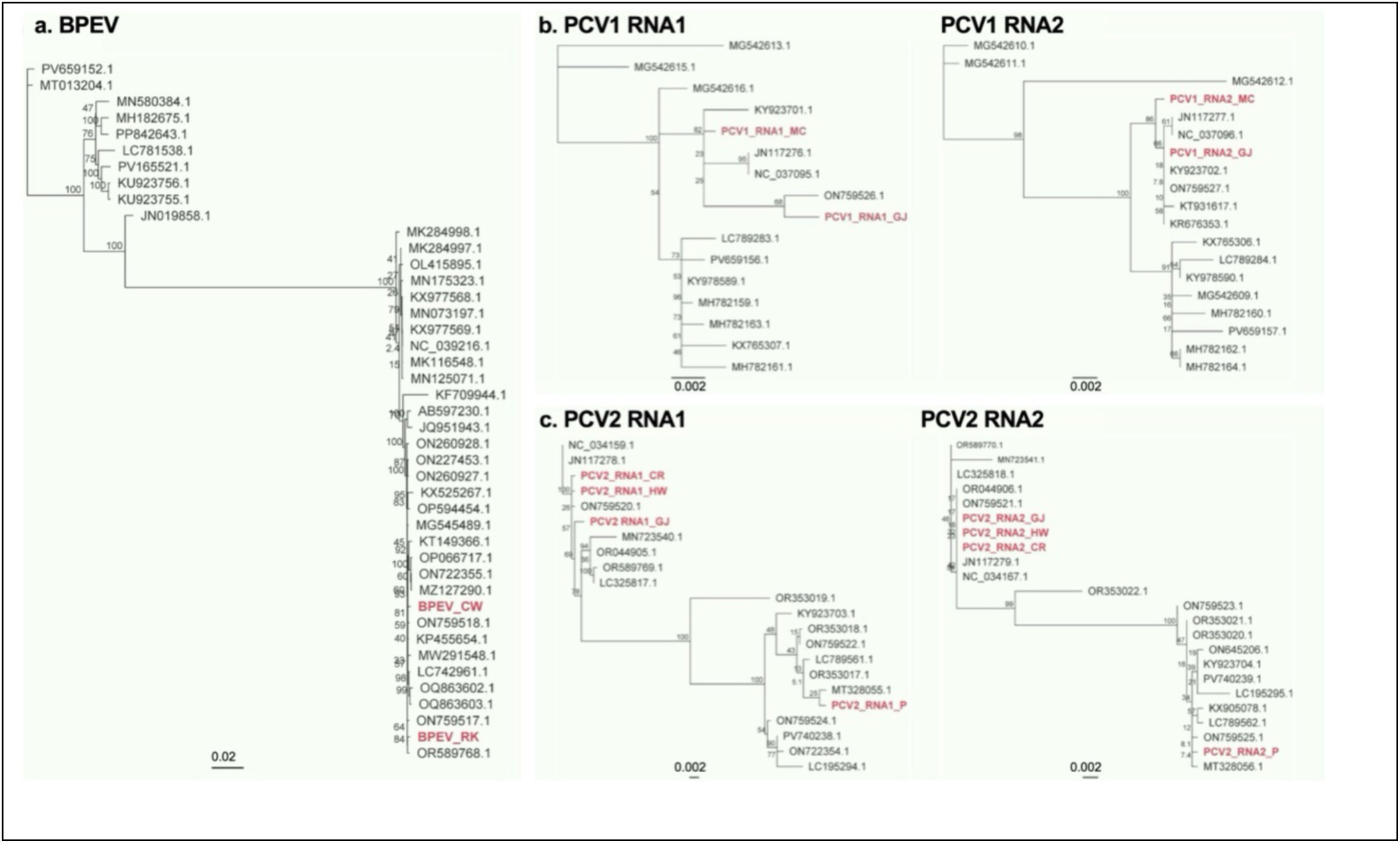
Phylogenetic trees for persistent viruses investigated in this study. Phylogenetic relationships based on full and near full-length nucleotide genome sequences of (a) bell pepper endornavirus (BPEV) sequences, (b) pepper cryptic virus 1 (PCV1) RNA1 and RNA2 sequences and (c) pepper cryptic virus 2 (PCV2) RNA1 and RNA2 sequences. Phylogenetic trees of viral nucleotide sequences were inferred in Geneious Prime using the PhyML Maximum Likelihood algorithm under the General Time Reversible (GTR) substitution model with gamma-distributed rate variation among sites. Bootstrap support was estimated using 1000 replicates, and values are shown. Branch lengths represent substitutions per site and persistent virus sequences from this study are highlighted in red bold type.

Plants of cultivars Gourmet Jalapeno, Mirasol Chili and Cayenne Long Slim carried PCV1 (Supplementary Table 3). Phylogenetic analysis of PCV1 sequences from Gourmet Jalapeno and Mirasol Chili revealed that these viruses clustered together for both RNA1 and RNA2 (Figure 1b and Supplementary Table 5). PCV2 was detected in four *C. annuum* cultivars (Gourmet Jalapeno, Poupila, Hungarian Wax and Serrano) and one *C. frutescens* line (Cayenne Red). PCV2 sequences from Hungarian Wax and Cayenne Red shared high identity across both RNA1 and RNA2. These sequences clustered separately from the PCV2 sequences from Poupila, indicating that the PCV2 variants of Hungarian Wax and Cayenne Red are distinct from PCV2 carried by Poupila (Figure 1c; Supplementary Table 5). The RNA 1 amplicon sequence for the PCV2 isolate identified in *C. annuum* cv. Serrano showed the closest identity with an isolate from the variety ‘Promotor’ (Tomašechová et al., 2020) (Supplementary Table 4), indicating that this isolate is also more similar to PCV2 from Hungarian Wax than to Poupila’s variant.

### Design and construction of vectors for virus-induced gene silencing of pepper persistent viruses and initial testing in *Nicotiana benthamiana*

Tobacco rattle virus (TRV) (Liu et al., 2002) was selected as a VIGS vector for two reasons. Firstly, TRV can initiate VIGS in pepper (Kim et al., 2017; Zhou et al., 2021). Secondly, TRV can induce VIGS against another virus. For example, TRV carrying a *green fluorescent protein* sequence inhibited infection of *Nicotiana benthamiana* plants by a potato virus X vector carrying the identical RNA sequence (Ratcliff et al., 1999). Since BPEV, PCV1 and PCV2 are all RNA viruses that replicate in the cytoplasm through the activity of their respective RdRps, the sequences selected as RNA silencing targets corresponded to those encoding the highly conserved GDD (gly-asp-asp) RdRp active site motif (Argos, 1988; Bruenn, 1991; Habili & Symons, 1989). Disrupting this sequence is highly effective in abolishing RNA virus replication (Olspert et al., 2015).

A plasmid carrying an infectious cDNA for TRV RNA2 (pYL156: Liu et al., 2002) was used to produce three constructs called pYL156:BPEV, pYL156:PCV1 and pYL156:PCV2 (Supplementary Figure 2a). These carried T-DNA sequences encoding recombinant TRV RNA2 with inserts corresponding to the respective GDD motifs and flanking residues within the BPEV polyprotein and the RNA1 open reading frames of PCV1 and PCV2 (Supplementary Figure 2b). Cells of *Agrobacterium tumefaciens* transformed with pYL156:BPEV, pYL156:PCV1 or pYL156:PCV2 were infiltrated into leaves of *N. benthamiana* plants together with cells carrying a plasmid encoding TRV RNA1 (pYL192: Liu et al., 2002). As a control for successful RNA silencing, agroinfection was also performed using cells carrying pYL192 (TRV RNA1) mixed with cells carrying a plasmid (pYL156:PDS) encoding a recombinant TRV RNA2 carrying a sequence designed to silence *phytoene desaturase* (*PDS*) transcripts (TRV-PDS) (Wang et el., 2022). RNA from systemically infected leaves was extracted at 12 days post-inoculation and used for RT-PCR analysis. Sequencing of TRV RNA2-derived PCR amplicons showed that inserts derived from *PDS* or persistent virus sequences were stable and had not undergone mutation or recombination events *in planta* (Supplementary Figure 3). These pilot agroinfection experiments using *N. benthamiana* showed that the recombinant TRV RNAs 2 encoded by pYL156:BPEV, pYL156:PCV1 and pYL156:PCV2 could be launched and replicate *in planta*, move systemically and remain genetically stable (Supplementary Figure 3).

### Pepper cotyledons are the optimal tissues for launching VIGS vectors by agroinfection

Several methods have been published for pepper agroinfection. Zhou et al. (2021) reported that launching of TRV-PDS infection in true leaves was aided by co-agroinfiltration with *A. tumefaciens* cells carrying plasmids encoding viral suppressors of RNA silencing (VSRs). However, with the pepper varieties and growth conditions used in this study, attempts to launch TRV-PDS by agroinfiltration of leaves with or without additional *A. tumefaciens* cells carrying T-DNA constructs for expression of either the cucumber mosaic virus 2b VSR or the tomato bushy stunt virus P19 VSR were unsuccessful. It appeared that *A. tumefaciens* induced necrosis true leaves, which prevented efficient launching of TRV-PDS (Supplementary Figure 4).

Consistent with two previous reports we found cotyledons to be the best sites for pepper agroinfection (Kim et al., 2017; Zhou et al., 2021). Optimisation experiments showed that neither the optical density of 600nm (OD_600_) for *A. tumefaciens* cell suspensions we used to agroinoculate *N. benthamiana* (OD_600_ = 0.5), nor the densities recommended previously by Zhou et al. (2021) (OD_600_ = 0.004) or Kim et al. (2017) (OD_600_ nm = 0.7) were sufficient to reliably launch TRV-PDS or silence *PDS* in pepper varieties used in this study. We found that infiltrating cells at an OD_600_ value of 2.0 for single or mixed cultures was consistently effective for successful agroinfection (Supplementary Figure 5).

### Persistent viruses can be silenced by VIGS in pepper plants and their progeny

We selected lines singly infected with either BPEV (Californian Wonder, Red King), PCV1 (Mirasol Chilli) or PCV2 (Hungarian, Poupila, Cayenne Red) for work to determine if VIGS was a viable approach for curing plants of persistent viruses. *A. tumefaciens* cells carrying pYL156:PDS were mixed with cells carrying constructs designed to silence BPEV, PCV1 or PCV2 RNAs (pYL156:BPEV, pYL156:PCV1 or pYL156:PCV2, respectively) and infiltrated into cotyledons. *In planta* TRV systemic movement is not uniform. Therefore, to visually track the progress of RNA silencing *A. tumefaciens* cells carrying pYL156:PDS were also included in agroinfection mixtures. This enabled photobleaching resulting from *PDS* silencing (Supplementary Figure 5) to be used both to confirm successful agroinfection as well as to guide tissue sampling of systemically infected tissue to monitor silencing of PCV1, PCV2 or BPEV RNAs using RT-PCR (Figure 2). The RNAs of BPEV, PCV1 and PCV2 all proved to be susceptible to VIGS. No evidence was found that TRV infection alone induced silencing of persistent viruses and when Gourmet Jalapeno plants were agroinfected using mixtures including cells carrying pYL156:PCV1, only PCV1 was silenced and PCV2 RNA levels were unaffected, demonstrating the specificity of the approach (Figure 2c).

**Figure 2.**
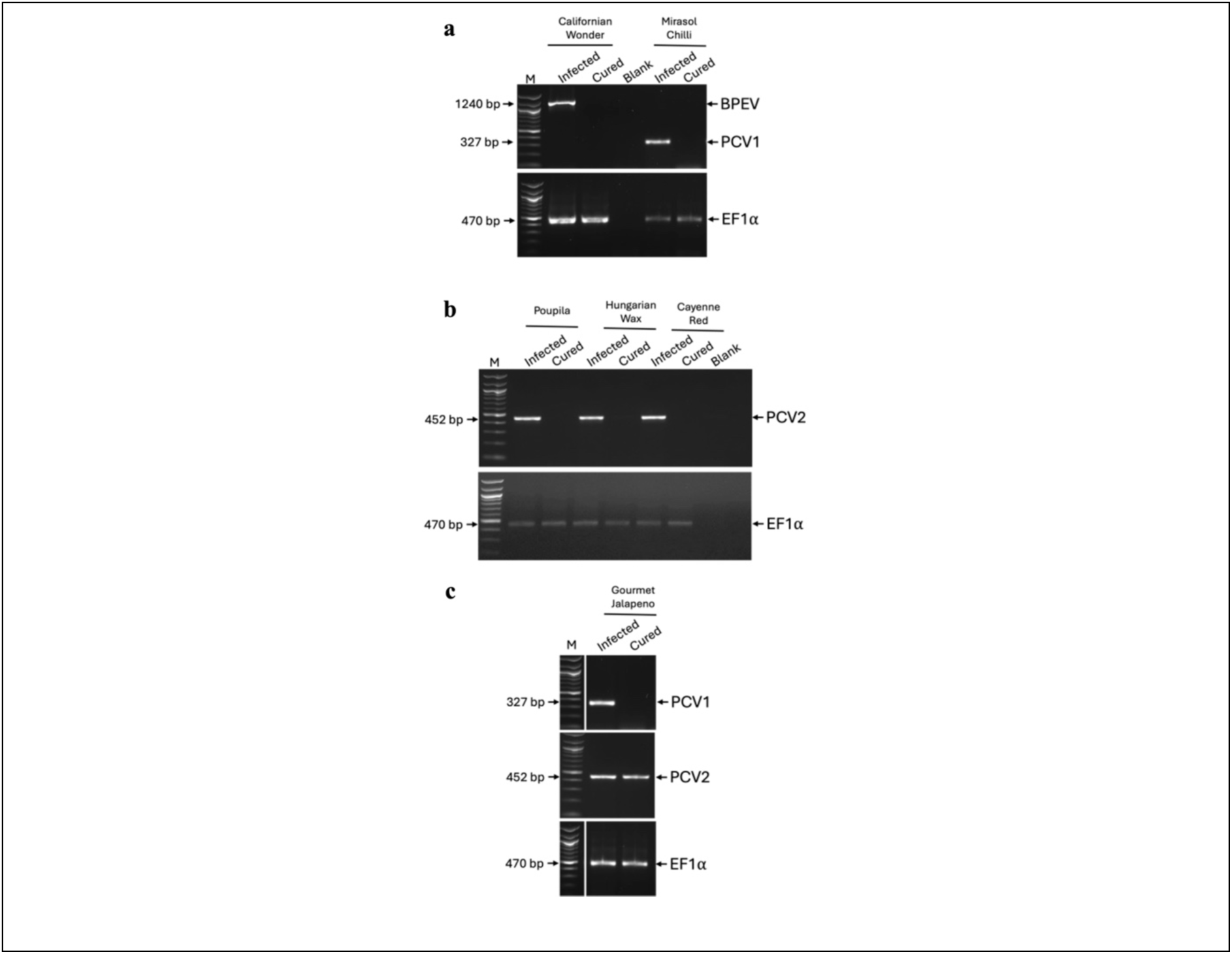
Silencing of persistent viruses in vegetative tissue of agroinfected pepper plants. RT-PCR using sequence-specific primers targeting PCV1, PCV2, and BPEV was performed on leaf samples from agroinfiltrated pepper plants. Products were resolved on 1% agarose gels. Californian Wonder and Mirasol Chilli cultivars subjected to triple agroinfiltration show examples of silenced and non-silenced lines for BPEV and PCV1 **(a)** and PCV2 **(b)**. In Gourmet Jalapeno, which is co-infected with PCV1 and PCV2, VIGS using pYL156:PCV1 successfully silenced PCV1 but left PCV2 unaffected **(c)**. Positive *CaEF1α* PCRs confirmed successful cDNA synthesis. ‘Blank’ indicates lanes loaded with no-template control reactions, and ‘M’ lanes loaded with a 100-bp DNA size marker ladder. Arrows indicate expected amplicon positions. Original gel images are shown in Supplementary Figure 11.

Silencing spread into flower peduncles and the fruits that developed subsequently (Supplementary Figure 6c, d). Photobleaching was used to guide selection of fruits for harvesting and collection of seed for subsequent analyses of the progeny of the agroinfected plants. The effectiveness of silencing of persistent virus RNAs varied markedly between cultivars (Supplementary Table 7). BPEV was silenced in one out of 43 agroinfected plants of the Californian Wonder cultivar, but in no Red King plants. PCV1 was silenced in 12 of 33 agroinfected Mirasol Chilli plants (Supplementary Table 7). Successful silencing PCV2 was variable in plants of the three cultivars carrying this virus (Poupila, Cayenne Red and Hungarian Wax) (Supplementary Table 7).

Seeds collected from plants that showed silencing of persistent viruses in the fruit tissue (Supplementary Figure 6D) were germinated and leaf samples were taken from the resulting plants for extraction of total RNA, which was used for RT-PCR to screen for the presence of persistent viruses and TRV. As expected, persistent viruses occurred in the progeny of control plants that had been agroinfected only with TRV-PDS (Figure 3). Furthermore, there were no instances where TRV or TRV-PDS was carried over to the progeny of agroinfected plants (Figure 3). Agroinfected plants in which a persistent virus was successfully silenced produced progeny that no longer carried (or were cured of) the persistent virus (Figure 3). Using seed from agroinfected plants in which persistent virus RNAs had or had not been silenced provided, respectively, cured and persistent virus-carrying isogenic lines for Californian Wonder (carrying or cured of BPEV), Mirasol Chilli (cured or carrying PCV1), and for Poupila, Cayenne Red and Hungarian Wax (retaining or cured of PCV2). For convenience we refer to these progeny plants as the ‘V1’ generation, i.e., the progeny of plants subjected to VIGS.

**Figure 3.**
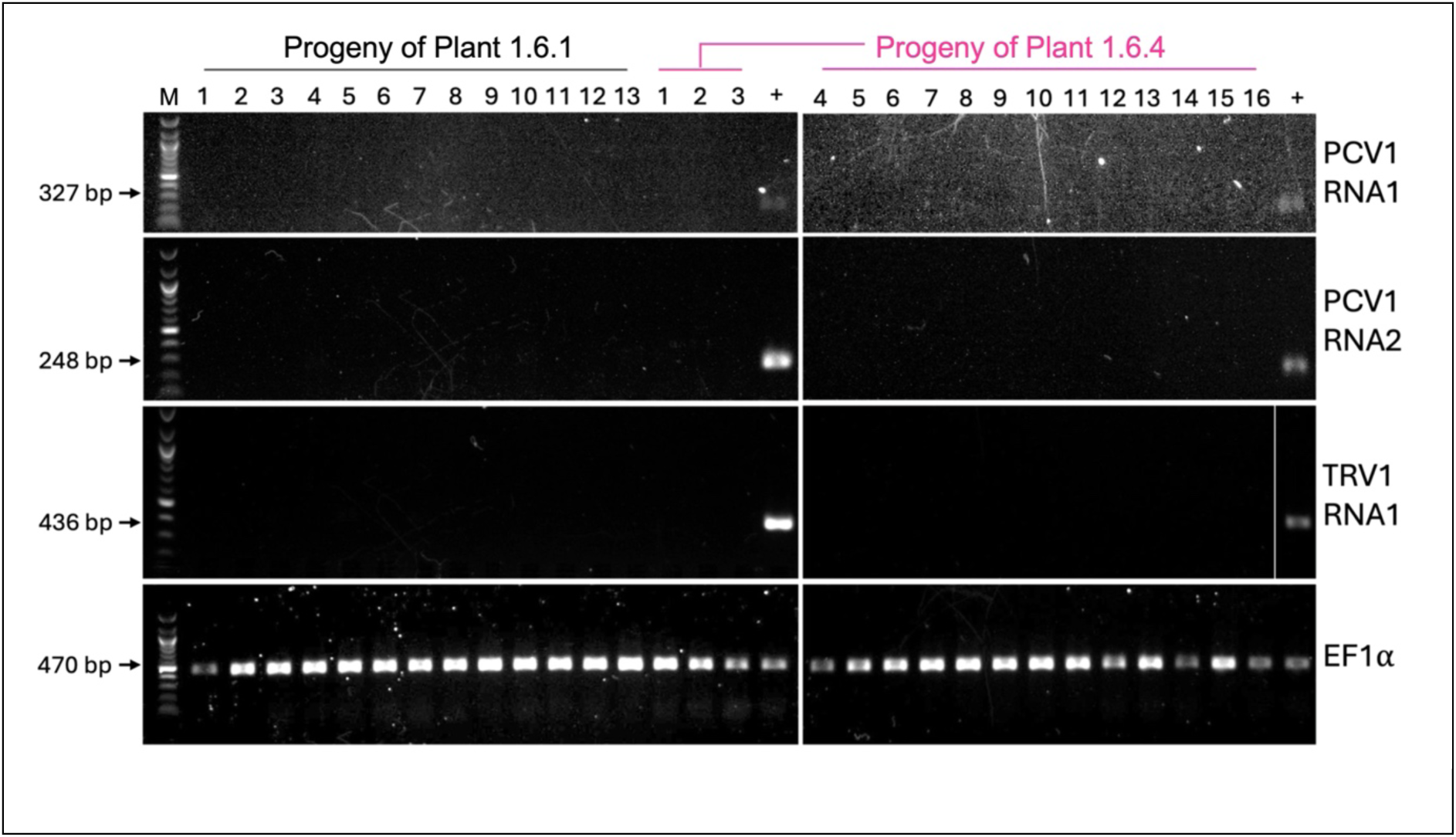
RT-PCR detection of viral RNAs in the progeny of silenced pepper plants. Reverse transcription PCR (RT-PCR) was performed using sequence-specific primers targeting PCV1 RNA1, PCV1 RNA2, and TRV RNA1 on tissue samples from progeny (i.e., V1 generation) of two Mirasol Chilli plants subjected to TRV-mediated VIGS (plants 1.6.1 and 1.6.4). Amplicons were resolved on 1% agarose gels. RNA from TRV-PDS agroinfected pepper plants served as a positive control (+) for persistent virus and TRV infection. Numbers below the cultivars designate individual screened plants. Results demonstrate successful co-silencing of PCV1 RNA1 and RNA2 in the progeny. TRV is not vertically transmitted to the progeny of agroinfected plants. RT-PCR amplification of *CaEF1α* served as an internal control, confirming successful cDNA synthesis across all samples. ‘Blank’ (B) contained no cDNA in the PCR reaction mixture. Lane ‘M’ was loaded with a 100 bp DNA marker ladder. Arrows indicate expected amplicon lengths for each RT-PCR target sequence. Original gel images are shown in Supplementary Figure 12.

### Persistent viruses can affect plant growth

The V1 progeny of agroinfected plants that had been cured or which still carried the persistent virus were grown, allowed to flower and seed harvested for further studies, and the characteristics of isogenic cured and uncured (control) plants were compared. We observed that persistent viruses can affect plant growth, with the most striking effects being observed in the Poupila background. From germination through 60 days post-germination, Poupila plants harbouring PCV2 were consistently and significantly taller than their cured counterparts with final heights approaching twice those of cured plants (Figure 4a). In contrast, PCV2 had no effect on the growth rate or final height of plants in the Cayenne Red background (Figure 4b). For Hungarian Wax, initial growth rates were similar between PCV2-harbouring and cured plants; however, Hungarian Wax plants carrying PCV2 reached significantly greater final heights than cured plants, being on average 19% taller (Figure 4c).

**Figure 4.**
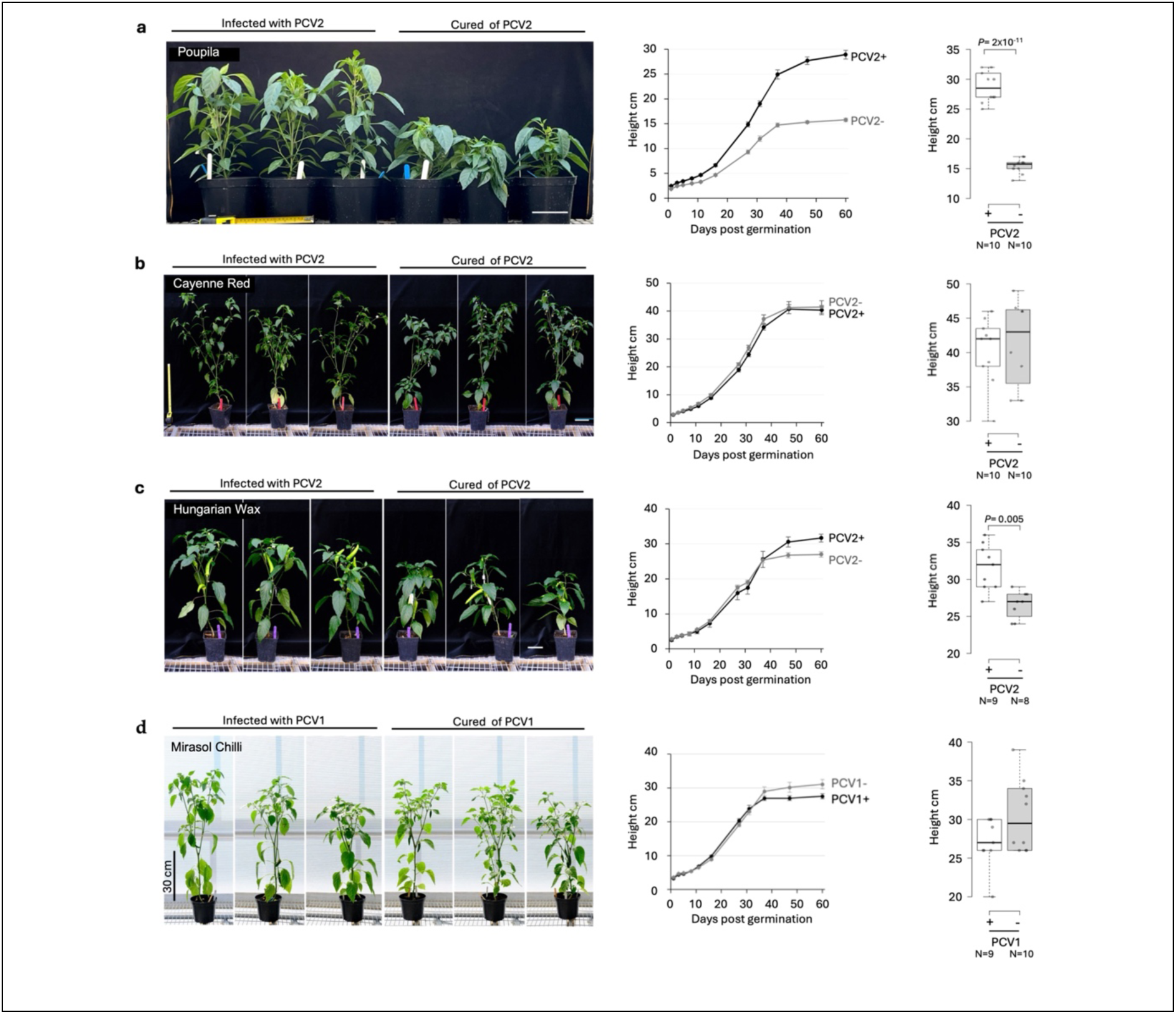
PCV2 enhances the growth of pepper plants in the Poupila background. Seeds of the V1 generation (i.e., the progeny of plants subjected to VIGS) from plants non-silenced or silenced for PCV1 (in the Mirasol Chili background) or PCV2 in three different genetic backgrounds (Poupila, Cayenne Red and Hungarian Wax) were germinated and subsequent plant growth observed. Plant height (measured to the apical bud or first branching point) was recorded at regular intervals (graphs) over a 60-day period (eventual heights shown in boxplots). Silencing of PCV2 in the Poupila background resulted in shorter plants (a), but not in Cayenne Red (b), while a small but statistically significant decrease in plant height occurred in cured plants in the Hungarian Wax background (c). Silencing of PCV1 in the Mirasol Chilli background had no significant effect on plant height (d). Scale bars in panels a–c: 10 cm.

Can PCV2 sequence differences explain the results? With respect to their RNA1 and RNA2 sequences, the PCV2 variants carried by Cayenne Red and Hungarian Wax plants are similar and occur in distinct clades from the corresponding sequences of the PCV2 of Poupila, indicating they are more closely related to each other than to the Poupila PCV2 variant (Figure 1). The predicted amino acid sequences of the RdRp open reading frames of the RNAs 1 of the Cayenne Red and Hungarian Wax PCV2 variants differed by only one amino acid (residue 95). In contrast, the RNA1 open reading frame of the Poupila PCV2 differed at 12 and 11 residues from the Cayenne Red and Hungarian Wax PCV2 RNA1 sequences, respectively (Supplementary Figure 7a). However, no amino acid differences occurred within known active sites or other conserved regions, e.g., the GDD domain (Supplementary Figure 7b). Furthermore, the predicted coat protein sequences encoded by the RNA2 open reading frames were identical for the PCV2 variants of Cayenne Red and Hungarian Wax, while both differed at the same five residues from the Poupila PCV2 coat protein sequence (Supplementary Figure 7c).

In the Mirasol Chilli background, plants cured of PCV1 showed no statistically significant differences in plant growth compared to plants carrying this partitivirus. Infected and cured plants grew at similar rates and by 60 days post-germination had reached comparable heights (Figure 4d). Californian Wonder background plants cured of BPEV were shorter than plants carrying BPEV at 35 days post-sowing, but this was not statistically significant. By 42 days post-sowing, cured and BPEV-infected plants showed no statistically significant difference in height (Supplementary Figure 8).

### Persistent viruses can influence flowering, fruit development and seed yield

Fruit was harvested at 120 days post-sowing from pepper plants still carrying PCV2 or cured of the virus (Figure 5). Fruit numbers on the day of harvest were similar for Poupila plants harbouring or cured of PCV2, though fruits from plants harbouring PCV2 appeared bigger with a larger proportion of ripe (i.e. red) fruit (Figure 5a). Poupila plants harbouring PCV2 or cured of this partitivirus began flowering at similar times (45 days post-sowing), suggesting that differences in fruit ripening were not due to delayed flowering in cured plants. The quantity of fruit was comparable between cured plants and those carrying PCV2 (Figure 5a, b). However, compared to cured plants, PCV2-infected plants produced significantly more seeds, with each fruit containing a greater mass of seeds (Figure 5c, e). This resulted in a significant overall improvement of ∼30% in total seed number and ∼32% total seed mass per plant (Fig. 5d, f).

**Figure 5.**
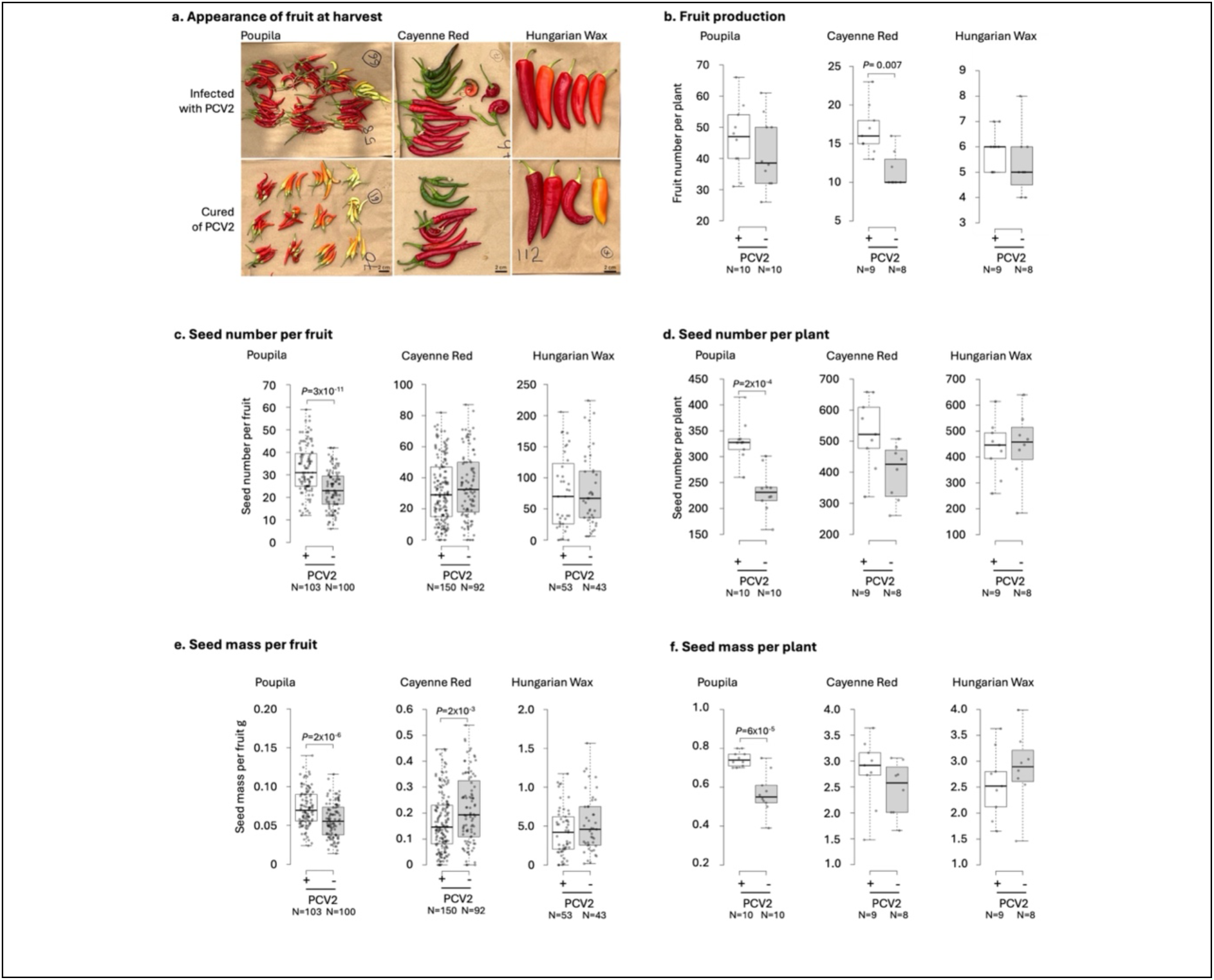
PCV2 enhanced seed production of plants in the Poupila background. Seeds from plants silenced for PCV2 (–) or still retaining the virus (+) in three backgrounds (Poupila, Cayenne Red and Hungarian Wax) were germinated and resulting plants were grown to maturity, allowed to flower and fruit was harvested (a). (b) With respect to fruit number, in the Cayenne Red background plants cured of PCV2 produced significantly fewer fruits than plants retaining PCV2; this was not seen in the other two backgrounds. In the Poupila background both seed number and seed mass per fruit were significantly lower in cured compared to PCV2-carrying plants, resulting in corresponding decreases in total seed number and seed mass per plant (c–f). The presence or absence of PCV2 did not significantly affect seed production in the Hungarian Wax background. The mass of seed produced per fruit was significantly greater in cured than in PCV2-carrying plants in the Cayenne Red background (e).

Cayenne Red plants harbouring PCV2 began flowering about one day earlier than their cured counterparts. However, PCV2-carrying plants produced significantly more fruit (∼46%; Figure 5a, b). Fruit from cured and PCV2-infected Cayenne Red plants contained a similar number of seeds, though PCV2 infection resulted in a lower aggregate seed mass per fruit (Figure 5c, e). Because PCV2-carrying plants produced more fruit and more seeds per plant than cured plants (Figure 5b, d), overall seed yield per individual was similar in both.

In the Hungarian Wax background, flowering times differed markedly between cured and PCV2-infected plants. Approximately half of the cured plants had begun to flower by 38 days after sowing and the rest by day 40. In contrast, only 50% of PCV2-carrying plants had flowered by day 45. However, flowering time differences were not associated with significant differences in fruit or seed numbers (Figure 5a–f).

In the Mirasol Chilli background, the presence of PCV1 was associated with significantly fewer fruit produced per plant (51% fewer) compared with cured plants (Figure 6a, b). PCV1-infected plants yield ∼25% fewer seeds per fruit than cured plants (Figure 6c). Overall seed production in terms of seed number and seed mass per plant in PCV1 harbouring plants was approximately half that of cured plants (Figure 6d, e).

**Figure 6.**
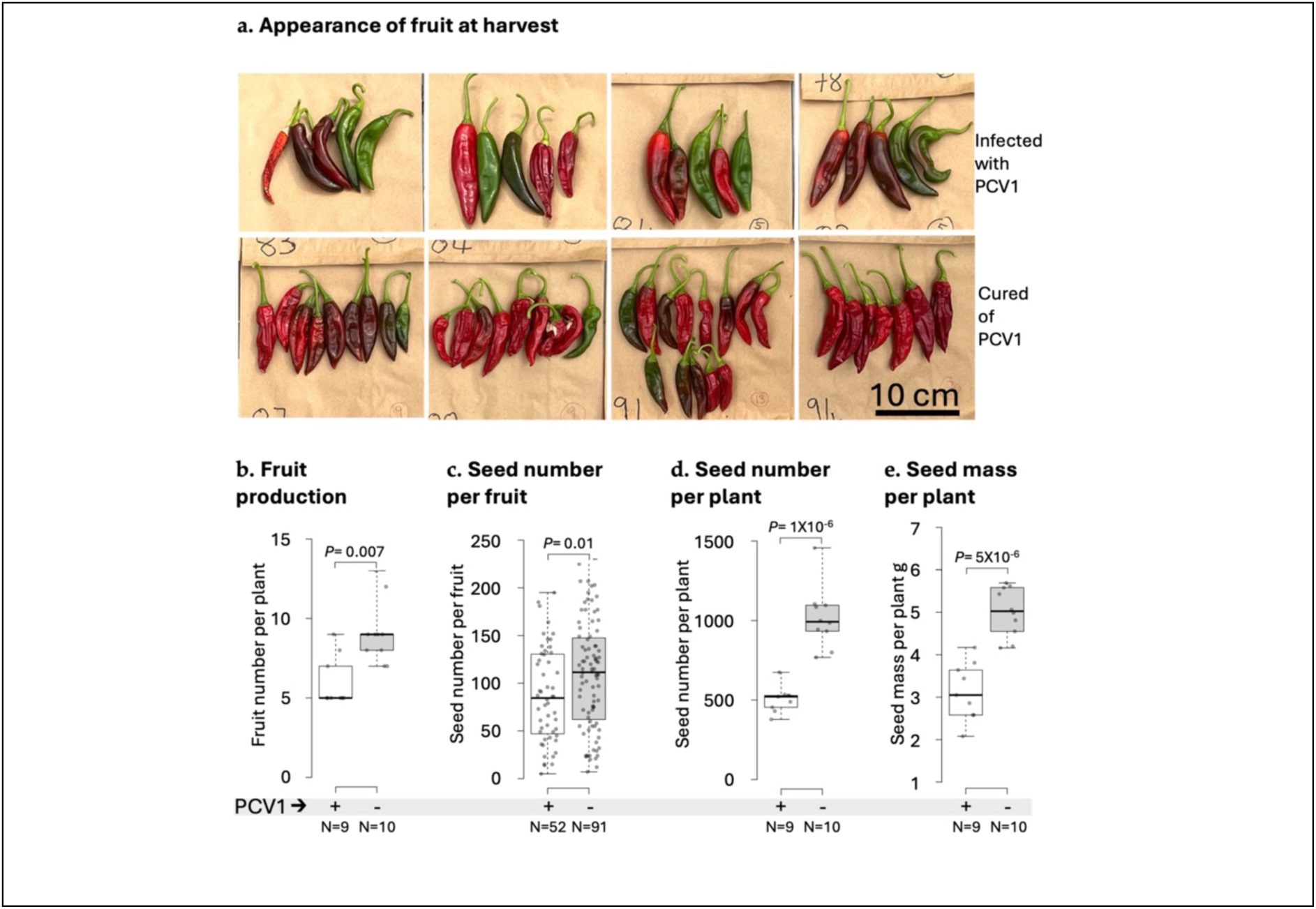
The effects of PCV1 on fruit and seed production. Seeds from plant lines cured of PCV1 (–) or still retaining PCV1 (+) in the Mirasol Chilli background were germinated and resulting plants were grown to maturity, allowed to flower and fruit was harvested. Fruit production (a, b), seed yield (c, d) and the overall mass of seeds produced (e) were all lower in PCV1-carrying plants than in their cured counterparts.

### Effects of persistent viruses on seed size and seedling development

To explore the effects of BPEV, PCV1 and PCV2 on seed size and the early post-germination plant development we used seeds of the ‘V2’ generation, defined here as the second generation descended from plants that had been subjected to VIGS, i.e., the seeds produced by V1 plants. It was found that both PCV1 and PCV2 could influence the size (quantified as cross-sectional area) and mass of individual seeds (Figure 7; Supplementary Table 8a, b). In plants harbouring PCV2 in the Hungarian Wax background seeds were significantly larger and heavier (12% and 7%, respectively) than those from cured plants. However, in the Cayenne Red background seeds produced by PCV2-carrying plants were significantly smaller and lighter than corresponding cured plants, while seeds from Poupila were similar in size and mass for cured and PCV2-carrying plants (Supplementary Table 8a, b). In the Mirasol Chilli background, plants harbouring PCV1 produced seeds that were ∼19% larger and ∼57% heavier than seeds produced by their cured counterparts; differences that were statistically significant (Figure 8a–c).

**Figure 7.**
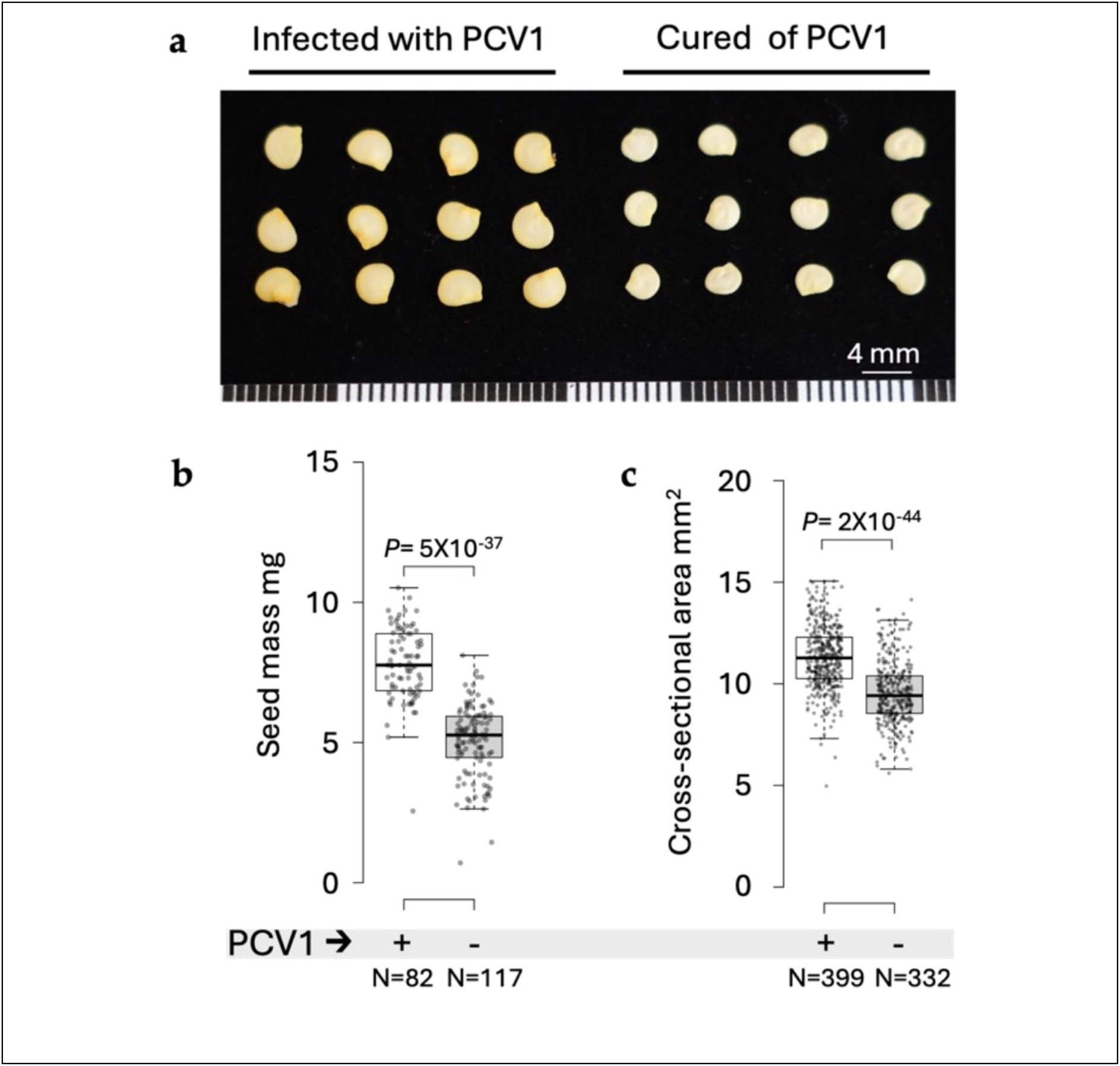
PCV1 enhances seed size in Mirasol Chilli. Seeds from plant lines cured of PCV1 (–) or still retaining PCV1 (+) in the Mirasol Chilli background were germinated and resulting plants were grown to maturity, allowed to flower, fruit was harvested and seeds recovered. These ‘V2’ seeds (i.e., the second generation after VIGS) were individually measured for mass and size (using seed cross-sectional area). (a) Representative seeds from PCV1-carrying and corresponding cured plants are shown. Seeds of PCV1-carrying plants had on average a mass 57% greater than seeds of cured plants (b), and their mean cross-sectional area was also significantly greater (c).

**Figure 8.**
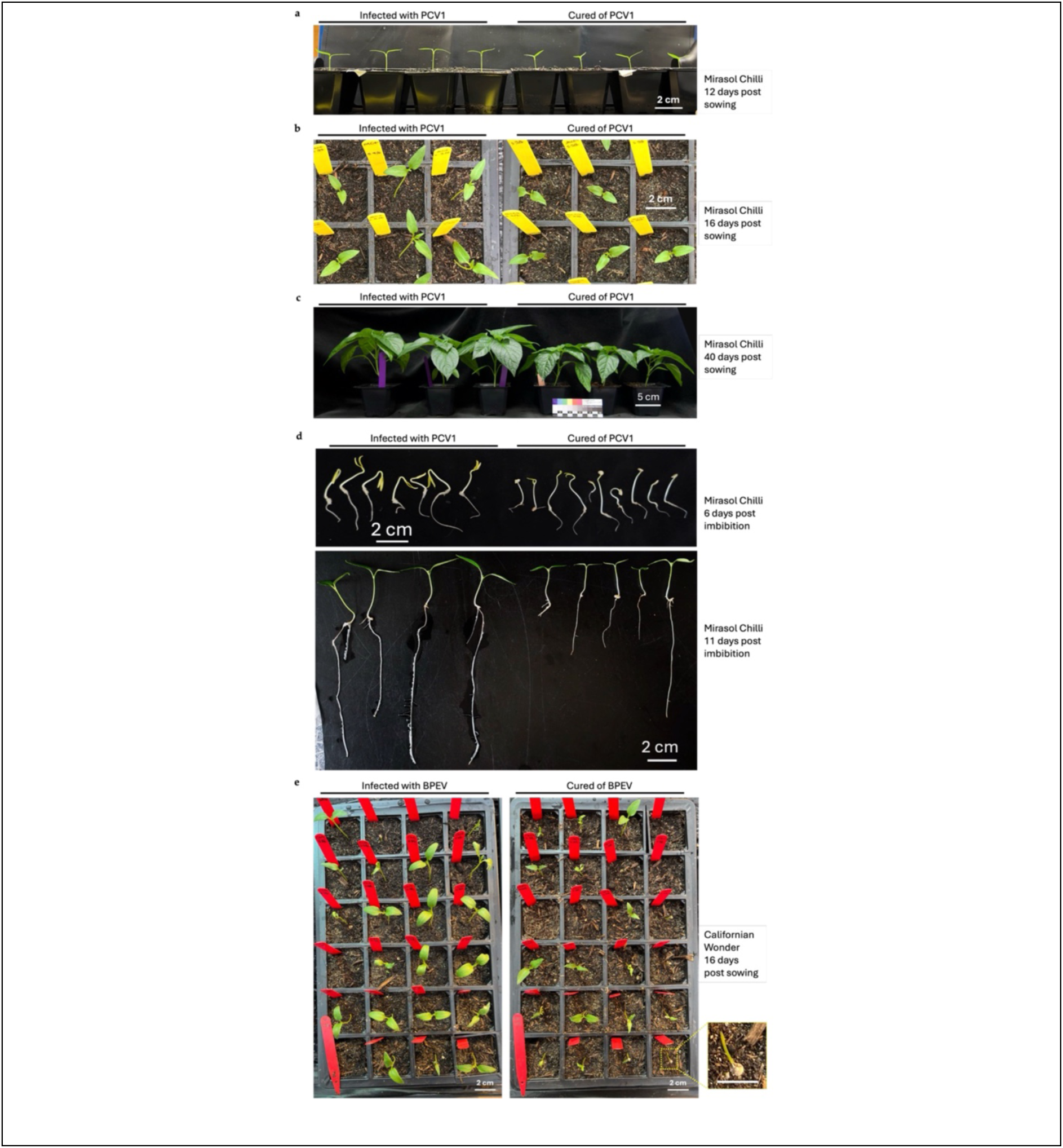
PCV1 and BPEV promote seedling vigour. Seeds from plant lines cured of PCV1 or BPEV or corresponding lines still retaining the viruses (Mirasol Chilli and Californian Wonder backgrounds, respectively) were germinated and the growth of the resulting seedlings was observed. In these experiments, seeds used were of the ‘V2’ generation (the second generation after VIGS, i.e., seeds produced by ‘V1’ plants, the progeny of plants subjected to VIGS). V2 PCV1-carrying seedlings developed faster than corresponding cured plants. They were markedly taller (a) and developed larger cotyledons (b), and this growth advantage persisted until at least 40 days after sowing (c). To assess root development, seeds were germinated on half-strength Murashige and Skoog agar and seedlings transferred to vertically suspended filter paper and photographed on a dark background. Root growth for PCV1-carrying seedlings was markedly faster than in corresponding cured seedlings (d). V2 seeds harvested from BPEV-carrying and corresponding cured Californian Wonder plants were sown directly into potting compost. Seedling establishment was less efficient for cured plants (e), with 11 of 36 seeds failing to germinate or emerge. A further five cured seeds germinated but remained seed-bound (i.e., cotyledons were trapped the seed coat: bottom right inset) and did not survive. In contrast, 20 of 36 BPEV-carrying seeds established successfully and seedlings produced fully expanded cotyledons.

BPEV-infected and cured Californian Wonder plants had, respectively, median fruit yields of 3 and 4 per plant, but this difference was not statistically significant (Supplementary Figure 9). We noted that in this bell pepper background the first fruit to develop tended to be largest and contained the most seeds, whilst many of the subsequently produced fruit were seedless. Resource competition between older and later-developing fruits on bell pepper plants may explain this (Marcelis & Baan Hofman-Eijer, 1997). Therefore, we recorded the total seed yield per plant rather than comparing seed number per fruit. Compared to cured plants, plants harbouring BPEV produced ∼24% more seeds with a total mass that was ∼25% greater, although these differences did not reach statistical significance (Supplementary Figure 9). BPEV-infected plants produced seeds that were moderately but significantly larger (∼6%) than cured plants (Supplementary Table 8; Supplementary Figure 9). Overall, there was no association between the presence or absence of the partitiviruses PCV1 and PCV2 with seed viability (within 6 months of harvesting), with infected and cured lines all exceeding ∼85% germination on MS agar and 70% on wet filter paper. Californian Wonder bell pepper plants harbouring or cured of BPEV had comparatively lower germination rates (60-70%) but were similar (Supplementary Table 9).

PCV1-carrying seed in the Mirasol Chilli background were larger than cured seed (Figure 7). Following germination this difference did not seem to have any marked influence on the timing of cotyledon emergence, but PCV1-carrying seedlings cotyledons were larger and the hypocotyls elongated more than for corresponding cured seedlings (Figure 8a, b). The growth advantage for V2 generation PCV1-carrying plants over their cured counterparts appeared to continue for at least 40 days after sowing (Figure 8c). Growing Mirasol Chilli hydroponically revealed that in addition to smaller cotyledons, the roots were smaller in cured seedlings than in seedlings retaining PCV1 (Figure 8d). BPEV had a more noticeable positive effect on seedling establishment (Figure 8e). Although the germination/seedling emergence rate was similar, cured plants were less uniform in size and many germlings failed to establish because of their inability to ‘push off’ the seed coat, resulting in death (see Figure 8d expanded inset showing detail of ‘seed-bound’ cotyledons).

For PCV2, there were variations in its effects across the three backgrounds Poupila, Hungarian Wax and Cayenne Red. Germination speed and cotyledon emergence/expansion occurred at similar rates for PCV2-infected and cured Cayenne Red seeds, while for Poupila cotyledon expansion was slower in cured plants but, contrastingly, cotyledon expansion progressed faster for cured Hungarian Wax seedlings (Supplementary Figure 10). The statistically significant effects of PCV1, PCV2 and BPEV on plant growth, fruit production and seed characteristics observed in this study have been summarised in Table 1.

**Table 1.**
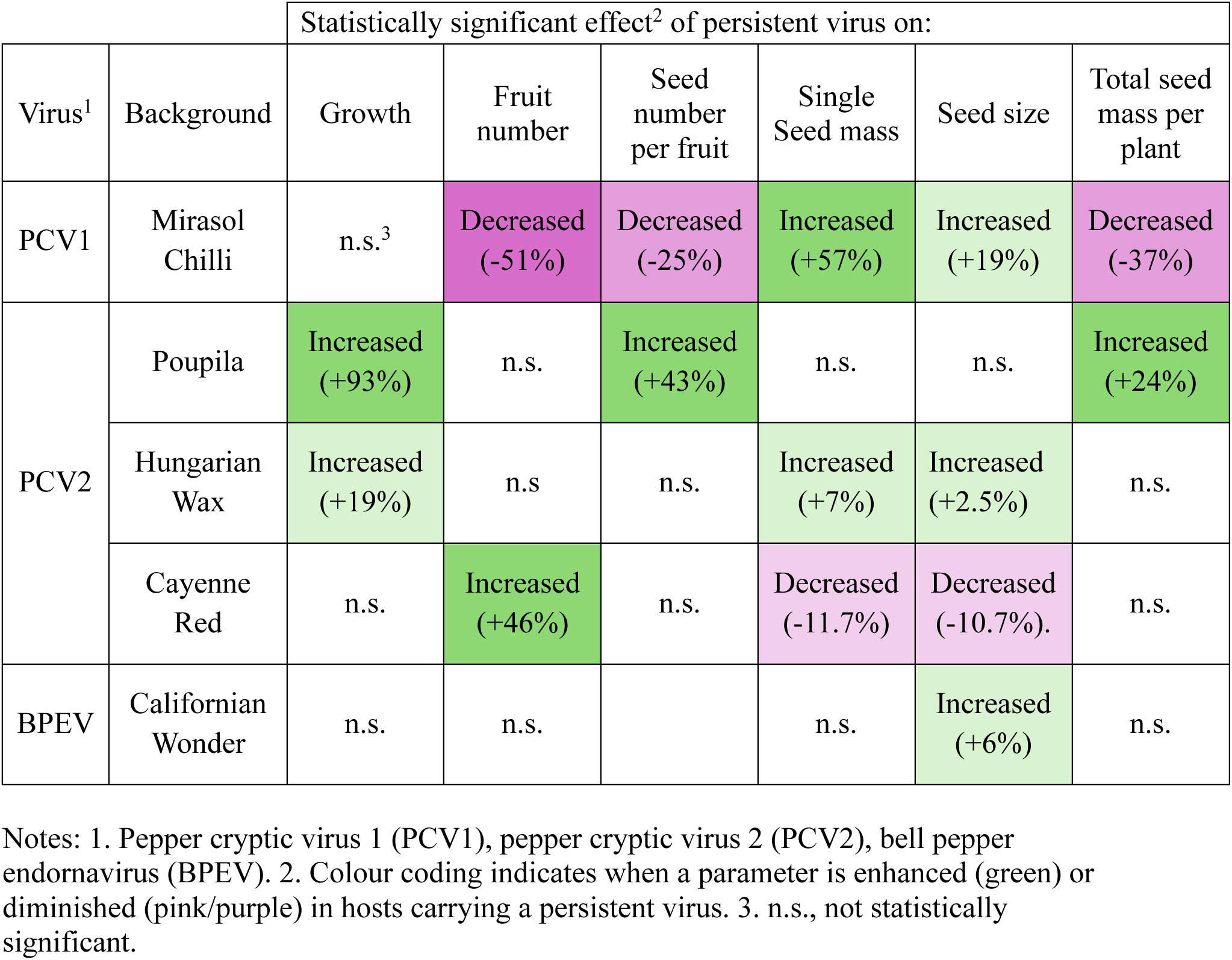
Summary of the effects of persistent viruses on host growth, fruit production and seed characteristics.

|  |  | Statistically significant effect<sup>2</sup> of persistent virus on: |  |  |  |  |  |
| --- | --- | --- | --- | --- | --- | --- | --- |
| Virus<sup>1</sup> | Background | Growth | Fruit number | Seed number per fruit | Single Seed mass | Seed size | Total seed mass per plant |
| PCV1 | Mirasol Chilli | n.s.<sup>3</sup> | Decreased (-51%) | Decreased (-25%) | Increased (+57%) | Increased (+19%) | Decreased (-37%) |
| PCV2 | Poupila | Increased (+93%) | n.s. | Increased (+43%) | n.s. | n.s. | Increased (+24%) |
|  | Hungarian Wax | Increased (+19%) | n.s. | n.s. | Increased (+7%) | Increased (+2.5%) | n.s. |
|  | Cayenne Red | n.s. | Increased (+46%) | n.s. | Decreased (-11.7%) | Decreased (-10.7%). | n.s. |
| BPEV | Californian Wonder | n.s. | n.s. |  | n.s. | Increased (+6%) | n.s. |

Notes: 1. Pepper cryptic virus 1 (PCV1), pepper cryptic virus 2 (PCV2), bell pepper endornavirus (BPEV). 2. Colour coding indicates when a parameter is enhanced (green) or diminished (pink/purple) in hosts carrying a persistent virus. 3. n.s., not statistically significant.

## DISCUSSION

The ubiquity of persistent viruses in many wild plants and their occurrence in lines of several crops is thought to indicate that they may confer benefits on plants (Roossinck 2010, 2013). It is plausible that under natural conditions carrying these viruses provides a selective advantage for wild plants, and that for domesticated plants lines bearing persistent viruses may have been unconsciously selected over many years by growers and crop breeders due to these viruses’ beneficial effects on agronomic characteristics (Roossinck, 2010; Fukuhara, 2019; Shates et al., 2019). Testing these ideas is important for two reasons. Firstly, to understand how persistent viruses might have shaped plant evolution under natural selection. Secondly, for practical considerations, i.e., to determine whether a persistent virus in a crop line is beneficial or deleterious, and to understand the potential of these agents as adjuncts to crop production or protection. This information could guide plant breeders as they develop new lines; for example, aiding in decisions for or against incorporating persistent viruses into breeding materials. Understanding the effects of persistent viruses requires a means of producing plants that are identical except for the presence of these agents. We have shown that VIGS provides a practical means of removing a persistent virus from a plant lineage to understand its effects on the host phenotype.

### Using VIGS to cure plants of persistent virus infection: Effectiveness and limitations

Nine pepper cultivars carrying PCV1, PCV2 or BPEV were identified. In some cases, for example Californian Wonder which harbours BPEV (Okada et al., 2011), persistent virus infection was already known, but novel examples were also identified. In six singly infected varieties the use of VIGS was successful and seed was recoverable from these plants for establishment of lines cured of infection with the endornavirus BPEV (Californian Wonder), or the partitiviruses PCV1 (Mirasol Chilli) and PCV2 (Hungarian Wax, Cayenne Red and Poupila). An additional set of experiments in which PCV1 was selectively silenced in the line Gourmet Jalapeno, which carries both PCV1 and PCV2, was carried out to show that VIGS-mediated curing was specific and not an artefact of using VIGS or a side effect of TRV infection.

In contrast to TRV-mediated transient silencing of *PDS* transcripts, the success of permanent silencing of persistent virus RNAs was variable, and in the present study silencing was not achieved for BPEV in the Red King background. This might reflect a limitation of TRV as a silencing vector in pepper and in future work using other viral vectors might improve success. However, it may simply be that while persistent virus RNAs replicate, while *PDS* transcripts cannot, the viral RNAs are harder to silence. Another reason may be that the relationship between the host RNA silencing system and persistent viruses is known to be complex. For example, knockdown of gene expression for a key host factor in antiviral silencing (Dicer-like 2) inhibited – rather than promoted – accumulation of Oryza sativa endornavirus RNA in rice (Urayama et al., 2010). If endornaviruses, and perhaps other persistent viruses, somehow depend on host RNA silencing systems to accumulate, this may have increased the difficulty of inducing RNA silencing to eradicate these viruses. In preliminary experiments we found that passaging TRV-PDS from agroinfected *N. benthamiana* to pepper plants using sap successfully induced silencing of pepper *PDS*. However, successful silencing of BPEV, PCV1 or PCV2 RNAs worked only when agroinfection was via the cotyledons, indicating that prolonged production of a silencing signal by a source tissue initiated at an early stage of plant development is required to clear these viruses from the host. A plausible explanation is that this is needed to disrupt the complex relationship (Urayama et al., 2010) between persistent virus RNA accumulation and host silencing.

### Persistent viruses affect seed and growth traits in pepper

Curing plants of persistent viruses altered one or more plant characteristics, indicating that although these agents do not induce disease symptoms as typically understood, i.e., stunting, deformation, chlorosis, mosaics etc., they do modify host phenotypes (Table 1). An important question is whether a persistent virus alters a host phenotype in a manner that increases host fitness, and by extension viral fitness since propagation of these viruses is inextricably linked with successful host reproduction. Seed properties perhaps offer an insight into this matter.

Curing Californian Wonder plants of BPEV decreased seed size and the overall mass of seed produced per plant. In the case of Mirasol Chili plants cured of PCV1, seeds were smaller and lighter, but cured plants produced more seed overall than PCV1-infected plants. PCV2’s influences on seed characteristics were investigated across three pepper backgrounds and distinct effects were noted. In the Poupila background a significantly higher seed number was associated with PCV2 infection, while PCV2 infection in both Poupila and Hungarian Wax was associated with larger and heavier seeds. For Cayenne Red, seed mass and size were similar for both cured plants and plants carrying PCV2.

It is generally thought that trade-offs exist for plants between producing fewer larger seeds with potentially higher seedling survival rates versus producing many smaller seeds with greater dispersal potential (Hodgson et al., 2020). But these trade-offs are not always clear cut and can be complicated further for animal-dispersed seeds (Coomes and Grubb, 2003); as would be the case for seeds of wild peppers (chiltipín, the ancestors of cultivated peppers), which are disseminated by birds (Tewksbury & Nabhan, 2001). The pepper-infecting persistent viruses studied here were associated with modification of seed numbers, seed size or both. If these effects occur in wild plants, they could be important for determining host competitiveness; potentially consistent with the hypothesis that persistent viruses are retained in plant lineages because they provide benefits to their hosts (Roossinck, 2012; Safari & Roossinck, 2018; Fukuhara, 2019; Takahashi et al., 2019). However, the plant lines used in this study are not wild plants, and in future work it would be interesting to further test this hypothesis by investigating the effects of curing plants of persistent virus-bearing chiltipín accessions on seed characteristics. As shown for BPEV the process of domestication of *Capsicum* species has itself influenced persistent virus evolution (Safari and Roossinck, 2018). Pepper plants that have been used, selected and distributed by humans are probably less strongly selected for seed size or number, although seed production can sometimes correlate positively with fruit yield (Marcelis & Baan Hofman-Eijer, 1997). Probably other traits with agronomic importance including some examined here (germination, growth, fruit production) as well as others not investigated in this study (e.g., stress resilience) have likely been unconsciously selected for. Using VIGS to investigate additional persistent virus-influenced phenotypes in a wider range of varieties will help elucidate the coevolution of persistent viruses with their hosts during domestication.

### The interplay of viral and host genotypes

The silencing of PCV2 in three plant backgrounds provided insights into the effects of viral versus host genotypes on persistent virus-host interactions. The RNA 1 and RNA2 sequences and the corresponding predicted protein sequences encoded by the respective open reading frames for the PCV2 strains present in Hungarian Wax and Cayenne Red were near-identical. But the PCV2 variants in Hungarian Wax and Cayenne Red are in a phylogenetic clade distinct from PCV2 carried by Poupila.

The PCV2 variant in Poupila was associated with greater plant height, higher seed production, size and mass, as well as increased fruit production. Fewer of these effects were apparent (at least to a statistically significant degree) in PCV2-carrying plants of the cultivars Cayenne Red and Hungarian Wax. This suggests that PCV2 strain differences explain some of the contrasting phenotypic effects of PCV2 infection in Poupila compared to the two other backgrounds. However, although Cayenne Red and Hungarian Wax carry nearly identical PCV2 isolates, different effects are associated with infection in each cultivar. For example, carrying PCV2 is associated with increased seed size in Hungarian Wax but not in Cayenne Red, and with increased fruit production in Cayenne Red but not in Hungarian Wax. A major difference between Cayenne Red and Hungarian Wax is that their species backgrounds are, respectively, *C. frutescens* and *C. annuum*. These comparisons show that the effects of PCV2, and by extension other persistent viruses, on plant phenotypes are influenced by both viral sequences and host genotype.

### Conclusions and limitations of the present study

Sequence-specific silencing of persistent virus RNAs using VIGS is an effective method for producing non-infected (cured) lines. Curing permits studies to be carried out on the effects of persistent viruses on plant phenotypes using non-infected and infected plants that are otherwise genetically identical. VIGS was successful in preventing transmission of persistent viruses of two distinct viral families. Indeed, it may also be a useful approach for clearing pathogenic seedborne vegetatively transmitted pathogenic viruses from plant material. The current study was limited to examination of the effects on plant growth and fruit and seed production caused by curing singly infected pepper lines. The partitiviruses PCV1 and PCV2 and the endornavirus BPEV had differing effects on these parameters, of which none appeared to be obviously deleterious to the host, as is more typically found with viral infections. Experiments with PCV2 in three host backgrounds indicate that viral sequences and host genotypes shape the outcome of the persistent virus–host relationship.

## MATERIALS AND METHODS

### Plant lines and growth conditions

Twenty pepper cultivars belonging to the species *Capsicum annuum* L., *Capsicum chinense* Jacq., *Capsicum baccatum* L., or *Capsicum frutescens* L. obtained from various sources listed in Supplementary Table 2 were screened for the presence of persistent viruses. Seeds were surface sterilised in 10% (v/v) sodium hypochlorite and 1% (w/v) sodium dodecyl sulphate for 10 min, before transfer to 70% (v/v) ethanol for 2 min, and rinsing in sterile water. Seeds were germinated in the dark on moist filter paper and seedlings transferred to high nutrient Levington Advance Pot & Bedding M3 compost mix (ICL, Ipswich, UK) in 15-cell growth tray with 6 x 6 cm cells (Berrycroft Stores Ltd., Willingham, UK). Plants grown for seed were transferred to 12 cm diameter 1.4 L pots. Seeds of pepper cultivars with germination rates below 40%, or which did not germinate within 10 d, were soaked in 200 mg.l^-1^ gibberellic acid (GA3) for 2 h to improve germination.

Seeds of *Nicotiana benthamiana* Domin. line RA-4 (Wylie et al., 2015) were germinated in 10 cm diameter plastic saucers containing Levington Advance F2 Seed & Modular Compost. Seedlings were transferred to 9 cm diameter pots containing a 50:50 mixture of M3 compost and F2 compost plus Miracle Gro continuous release plant food (Evergreen Garden Care Ltd, Surrey, UK). Pepper and *N. benthamiana* plants were cultivated in controlled environment rooms (Conviron, Manitoba, Canada) maintained at 22°C and 60% relative humidity, under a 16 h photoperiod with 200 μE.m^-2^.s^-1^ of photosynthetically active radiation.

### Detection and quantification of persistent virus and tobacco rattle virus RNA

To detect viral RNAs, total RNA was extracted from plant tissue for reverse transcription-coupled polymerase chain reaction (RT-PCR) assays and for reverse transcription-coupled quantitative polymerase chain reaction (RT-qPCR) assays as recently described (Brine et al., 2023a,b) using appropriate primers (Supplementary Table 1). Amplicons were subjected to automated Sanger DNA sequencing (Sanger et al., 1977; Smith et al., 1986). Methods for metatranscriptome library construction, high-throughput sequencing, non-host unigene assembly and phylogenetic analyses are described in Supplementary Methods.

### Inoculation with TRV

Plasmids carrying inserts for infectious clones of TRV RNA 1 (pYL196) and a modified TRV RNA 2 (pYL156) (Bachan and Dinesh-Kumar, 2012; Liu et al., 2002) were obtained from Addgene (https://www.addgene.org/ accessed November 12, 2024). The plasmid pYL156:PDS encoding an infectious TRV RNA 2 clone carrying a tomato-derived sequence designed to silence the *PDS* transcript (Wang et al., 2022) was a generous gift of Dr Zhengming Wang (Department of Plant Sciences, University of Cambridge). The plasmids pYL156:BPEV, pYL156:PCV1 and pYL156:PCV2 were produced by insertion of sequences corresponding to the regions of the BPEV ORF and of the RNAs 1 of PCV1 and PCV2 encoding the conserved GDD motif and flanking amino acid sequences of these viruses’ respective viral RdRp enzymes (Supplementary Figure 2). Plasmids were transformed into cells of *Agrobacterium tumefaciens* GV3101 for use in agroinfection experiments.

Three days before an agroinfiltration or agroinfection experiment, *A. tumefaciens* GV3101 cells containing the appropriate plasmids were streaked on LB (Sambrook et al., 1989) agar plates with 50 μg.ml^-1^ kanamycin and 10 μg.ml^-1^ rifampicin. After 48 h at 28°C, single colonies of *A. tumefaciens* containing the desired construct were inoculated to 10ml liquid cultures (LB containing 50 μg.ml^-1^ kanamycin and 10 μg.ml^-1^ rifampicin) for 12 h at 28°C with aeration at 200 rpm before cells were collected by centrifugation at 4,500 x *g* for 15 min. Pelleted cells were resuspended in freshly made agroinfiltration solution (10 mM MgCl_2_, 10 mM 2-morpholinoethanesulfonic acid pH 5.6 and 150 μM acetosyringone) and incubated for 2 h at room temperature (Watt et al., 2019). For agroinfiltration of *N. benthamiana* plants, cell suspensions were adjusted to an OD^600^ of 0.5 before introducing the cells, using a needle-less syringe, into the abaxial surfaces of leaves of 3-week-old plants (Watt et al., 2019). However, for efficient agroinfection of pepper plants, suspensions of *A. tumefaciens* cells were adjusted to an OD^600^ of 2.0 and infiltrated into the cotyledons of seedlings before expansion of true leaves. Cultures of *A. tumefaciens* cultures carrying different plasmids were combined as needed using equal volumes of resuspended cells. Passaging of modified TRV from systemically infected *N. benthamiana* plants to pepper by mechanical inoculation was carried out by homogenising 2–3 systemically infected leaves in 0.01M potassium phosphate pH 7.0 and applying the homogenate with a cotton bud to cotyledons sprinkled with Carborundum (320 grit silicon carbide powder: Alfa Aesar, Thermo-Fisher Scientific, UK). Mock inoculation used buffer only. Following inoculation, cotyledons were rinsed with sterile water and plants covered for 24 h to maintain humidity.

### Plant growth, fruit and seed measurements

Seeds harvested from plants subjected to VIGS i.e., ‘V1’ generation, were germinated and plants cultivated as described above in the University of Cambridge Botanic Garden glasshouse maintained at 18–25°C with approximate humidity of 55%. During periods of the year when daylight levels fell below 150 W/m^2^ additional lighting (Lucalux LU 400 W/PSL) automatically activated between the hours of 0400 and 2000. Virus infection status of plants was confirmed by preparing leaf RNA samples for analysis by RT-PCR.

Plant heights were recorded for up to 60 days, measuring from the compost line to depending on cultivar the apical bud or to the first branching point once branching occurred. For Mirasol Chilli, Cayenne Red, Hungarian Wax and Poupila cultivars fruit was harvested at 120 days post-sowing. For the Californian Wonder cultivar fruit was harvested 130 days post sowing. Fruit harvested from individual plants was stored in paper bags and sorted by approximate size and ripeness. Seeds were extricated and air-dried in Petri dishes over 14 days. Each seed batch was labelled to maintain traceability to its specific fruit number and parent plant. While seeds dried, they were counted with the aid of a tally counter to obtain figures for yield. Due to the high productivity of Mirasol Chilli, seed sampling was standardized by extracting seeds exclusively from the first 10 mature, ripe fruits produced per parental line.

Dried seed batches (per fruit) were weighed and transferred to 7x10cm paper bags and stored at room temperature. To evaluate individual seed masses, 12 seeds from a single fruit were weighed individually using a microbalance (MX5, Mettler-Toledo Inc., Columbus, OH, USA).

Major cross-sectional area was used a proxy for seed size. Between 20–100 seeds from single fruits were photographed on a black background. Using ImageJ software (v1.54, NIH, USA), digital images were converted to 8-bit grayscale, thresholded to isolate individual seeds from the background and the “Analyze Particles” function applied. A physical scale was included in each image to enable spatial calibration.

### Analysis of seed viability and seedling vigour

The germination rate of V2 generation seed (i.e., the second generation after VIGS) was done by two methods. Seeds were either germinated in the dark on moist filter paper as described above or were surface-sterilized and placed on half-strength Murashige and Skoog (MS) basal medium (Murashige and Skoog, 1962) solidified with agar and incubated at 28 °C in the dark for 12 days. Germination was scored as successful once the radicle emerged from the seed coat. To assess seed vigour, seeds were sown directly into a moistened 4:1 (v:v) substrate mixture of Levington Advance Pot & Bedding M3 compost with horticultural sand in P24 potting tray inserts (individual cell volume dimensions of 50 x 48 x 50 mm: w x l x h) (Berrycroft). Because subterranean germination could not be monitored directly, success was evaluated based on the emergence of germinated seedlings and subsequent cotyledon expansion.

### Statistical approaches

Data formatting and descriptive statistics were performed using Microsoft Excel for Mac (Version 16.89.1, Microsoft Corporation, USA). To evaluate differences between specific treatment pairs (i.e., plants infected with or cured of persistent viruses), two-tailed, independent Student’s *t*-tests were performed. To control the family-wise error rate across multiple comparisons, critical *p*-values were adjusted using the Bonferroni correction. Differences were considered statistically significant when the calculated probability fell below the corrected threshold at an adjusted α = 0.01. Boxplots were generated using ‘BoxPlotR: a web-tool for generation of box plots’ available at http://shiny.chemgrid.org/boxplotr/ (accessed September 5 2026) (Spitzer et al., 2014). Centre lines represent medians; box limits indicate the 25th and 75th percentiles (calculated using R). Whiskers extend to minimum and maximum values for sample sizes under 100, or to the 5th and 95th percentiles for sample sizes exceeding 100.

## ACKNOWLEDGMENTS

We are grateful to Francis O. Wamonje, Tom Brine and Sam Crawshaw for useful discussions, Dr Zhengming Wang for the TRV-PDS construct, and the Cambridge University Botanic Garden for gifts of certain *Capsicum* lines (Supplementary Table 2). We are grateful for major support from The Leverhulme Trust (RPG-2022–134) and the UK Biotechnological and Biological Sciences Research Council (UKRI2973). SBV received supported from a Cambridge Trust PhD scholarship.

## AUTHOR CONTRIBUTIONS

JPC and AMM conceptualized and designed the study. SBV, AMM, AEP, SLM and LME carried out the experimental work. AMM, SBV and JPC analysed the data. JPC, AMM and SBV drafted the manuscript. All authors revised and approved the manuscript.

## SUPPORTING INFORMATION

Additional Supporting Information may be found in the online version of this article.

## DATA AVAILABILITY STATEMENT

The authors declare that the main data supporting the findings of this study are available within the article and its Supplementary Information files, methods, and materials. Sequencing, metagenome and whole-genome sequencing data can be accessed at: https://eur03.safelinks.protection.outlook.com/?url=https%3A%2F%2Fwww.ncbi.nlm.nih.gov%2Fbioproject%2F%3Fterm%3DPRJNA1271429&data=05%7C02%7Cjpc1005%40cam.ac.uk%7C995f497322324376687208df0b5652e1%7C49a50445bdfa4b79ade3547b4f3986e9%7C1%7C0%7C639242137707900965%7CUnknown%7CTWFpbGZsb3d8eyJFbXB0eU1hcGkiOnRydWUsIlYiOiIwLjAuMDAwMCIsIlAiOiJXaW4zMiIsIkFOIjoiTWFpbCIsIldUIjoyfQ%3D%3D%7C0%7C%7C%7C&sdata=Hv%2BV%2BHcWotOpZ%2FcyUn7FOWmmIMVmqc7NtDGwyiILuVk%3D&reserved=0 (Accessed September 5 2026).

## RESOURCE AVAILABILITY

Requests for further information, resources and materials should be directed to, and reasonable requests will be fulfilled, by the lead contact: John P. Carr.

## Supplementary Figures and Materials and Methods

### Supplementary Figures

**Supplementary Figure 1.**
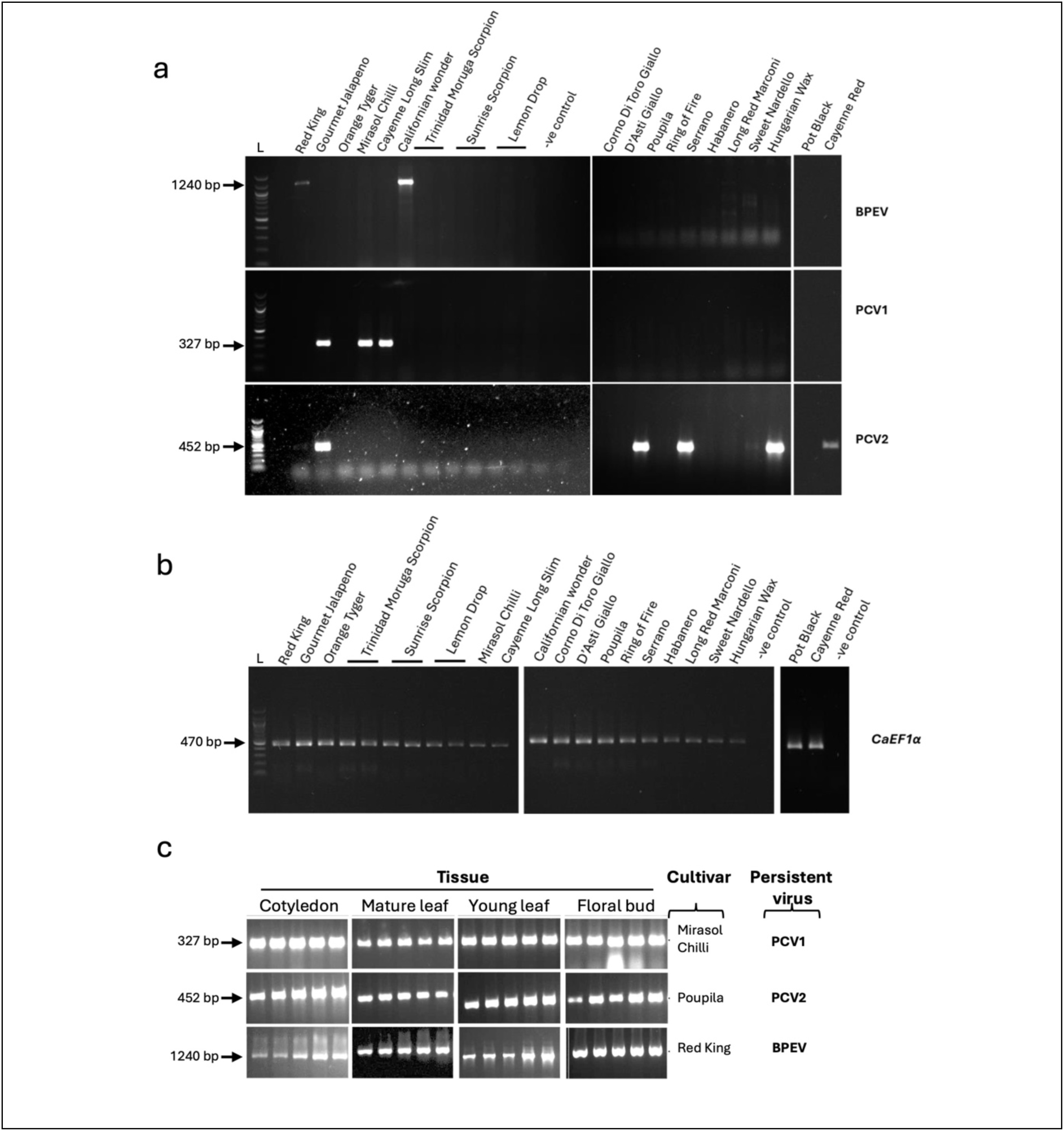
Screening for PCV1, PCV2 and BPEV RNAs and accumulation in different plant tissues. RT-PCR detection of BPEV, PCV1, PCV2 (a), and the pepper reference gene *CaEF1α* (b) in 20 pepper cultivars. PCR products were resolved on 1% agarose gels. Arrows indicate detected persistent virus amplicons. *CaEF1α* amplification confirmed successful cDNA synthesis. L, 100-bp DNA ladder; –ve, no-template control. (c) RT-PCR for PCV1, PCV2 and BPEV was performed on RNA extracted from cotyledons, mature and young leaves, and floral buds of five biological replicates from cultivars hosting the respective viruses: Mirasol Chilli (PCV1), Poupila (PCV2), and Red King (BPEV). Amplicon sizes are shown in base pairs (bp).

**Supplementary Figure 2.**
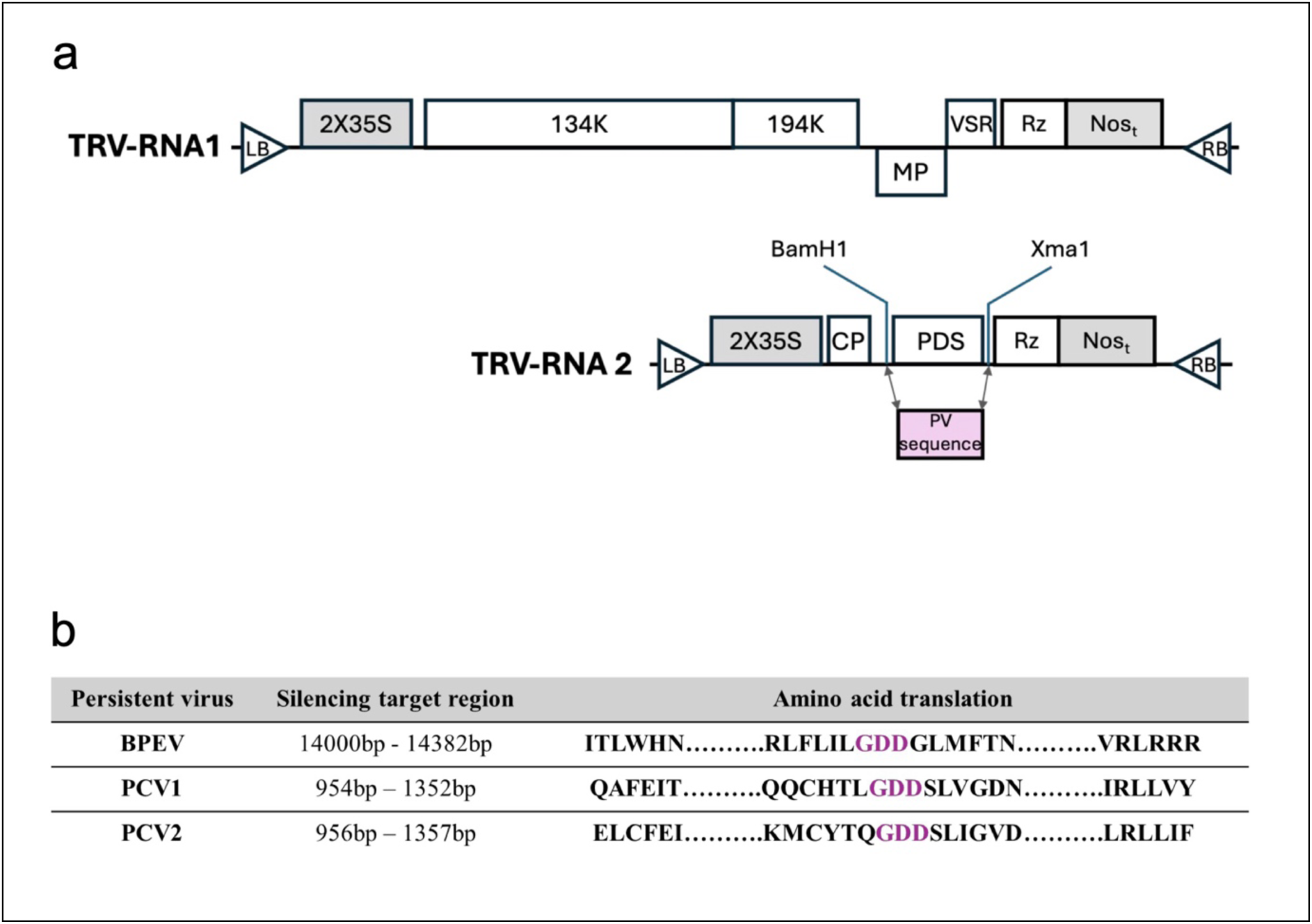
TRV based VIGS vectors (after Bachan and Dinesh-Kumar, 2012). TRV cDNA clones had been placed in-between the duplicated CaMV 35S promoter (2×35S) and the nopaline synthase terminator (NOSt) in a T-DNA vector. LB and RB refer to left and right borders of T-DNA. Rz, self-cleaving ribozyme. BamH1 and Xma1 restriction enzymes were used to remove the phytoene desaturase (PDS) sequence and replace with the sequences homologous to the respective GDD motifs (and adjacent sequences) present in the BPEV polyprotein and the RNA1 open reading frames of PCV1 and PCV2 (pYL156:BPEV, pYL156:PCV1 and pYL156:PCV2, respectively). Sequences for VIGS constructs were: BPEV RdRp from California Wonder, PCV1 RdRp segment from Gourmet Jalapeno and PCV2 RdRp from Poupila (b).

**Supplementary Figure 3.**
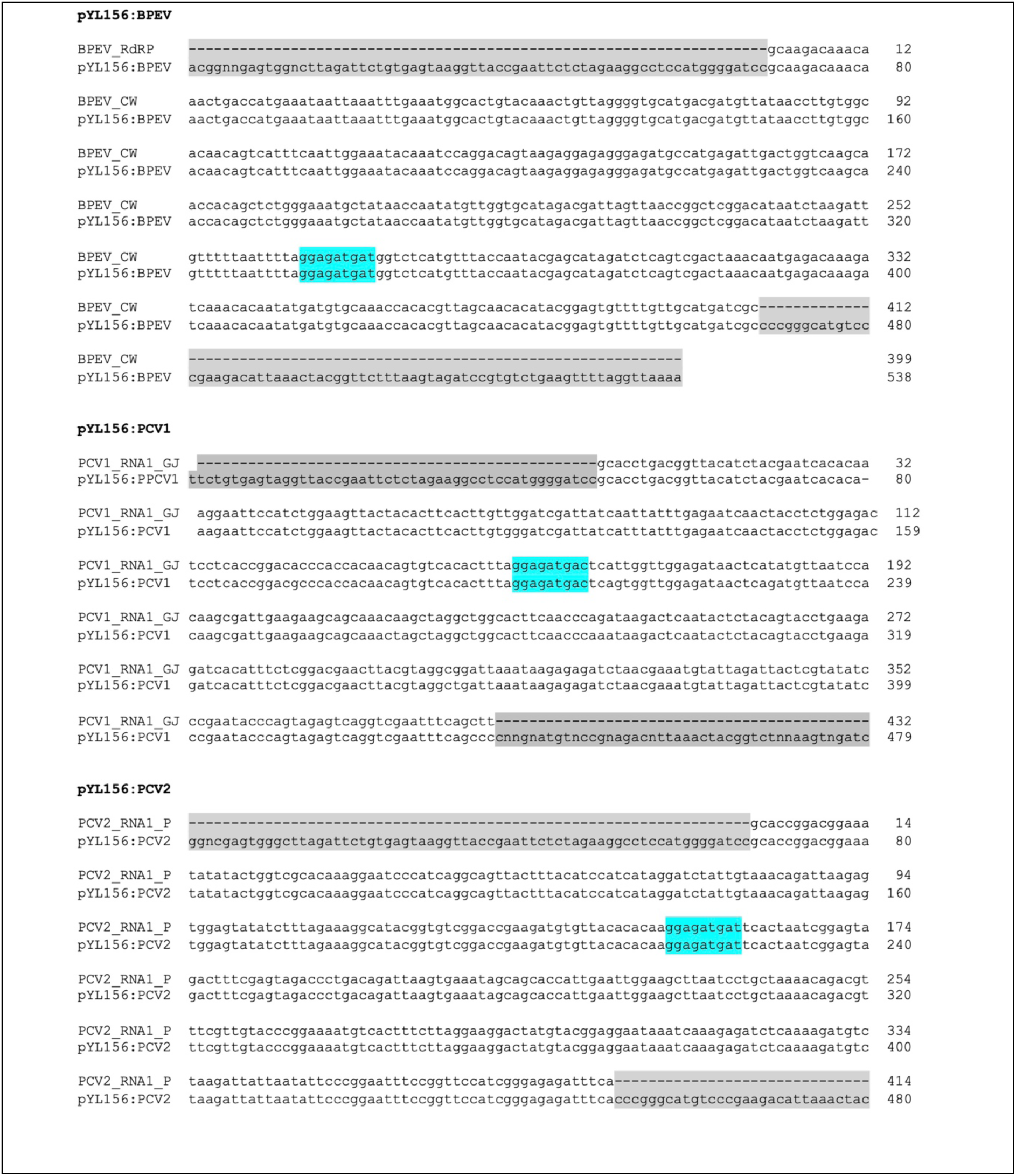
TRV-RNA2 VIGS constructs remained stable *in planta.* Viral RNA was recovered from systemically infected *N. benthamiana* leaves and RT-PCR followed by sequencing of the recombinant TRV RNAs 2 confirmed that the inserted sequences remained free of mutations or recombination. Sequence alignments show the original construct sequence (bottom row) aligned with the RT-PCR product (top row) from each infected plant. TRV-derived regions are shaded in grey; nucleotides encoding the conserved GDD motif within each persistent virus RdRP is highlighted in blue. RdRP sequences derived from the VIGS-modified TRV RNA isolated from systemically infected leaves exhibited 100% sequence identity with their respective constructs in the case of BPEV and PCV2, and 96.9% similarity in the case of PCV1.

**Supplementary Figure 4.**
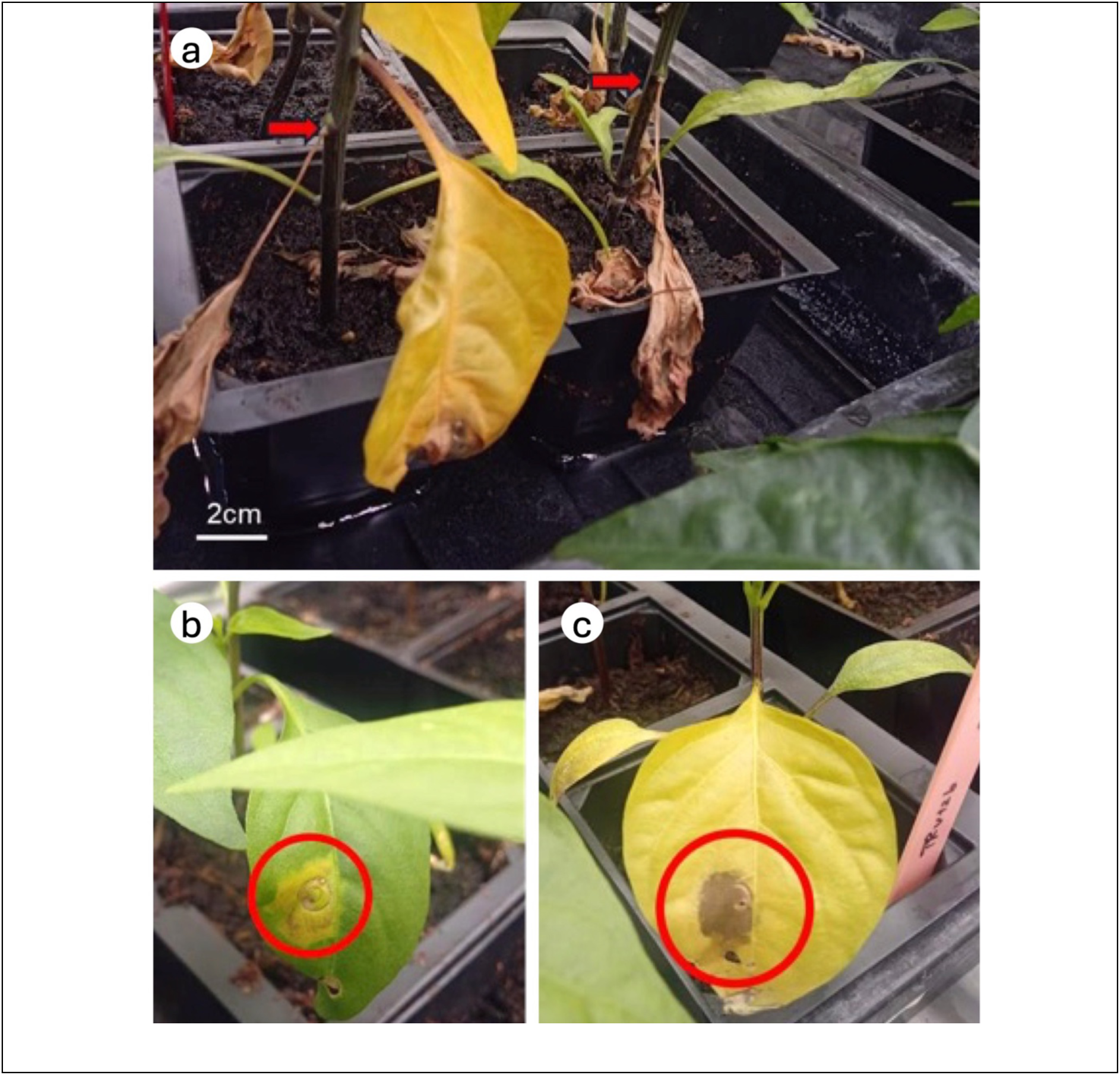
Examples of leaves of pepper plants agroinfiltrated with TRV1 + TRV2:PDS (without or with viral suppressors of RNA silencing). (a) Pepper plants agroinfiltrated with TRV1 + TRV2:PDS did not exhibit *PDS* silencing symptoms. By two weeks post-agroinfiltration, the infiltrated leaves had died (red arrows). Co-agroinfiltration with TRV1 + TRV2:PDS and either pBIN61:p19 (b) or pSITE:CMV2b (c). Neither RNA silencing suppressor enhanced TRV infection efficiency and there was no induction of *PDS* silencing. Necrosis developed in the infiltrated leaf regions (red circles) within two weeks of agroinfiltration.

**Supplementary Figure 5.**
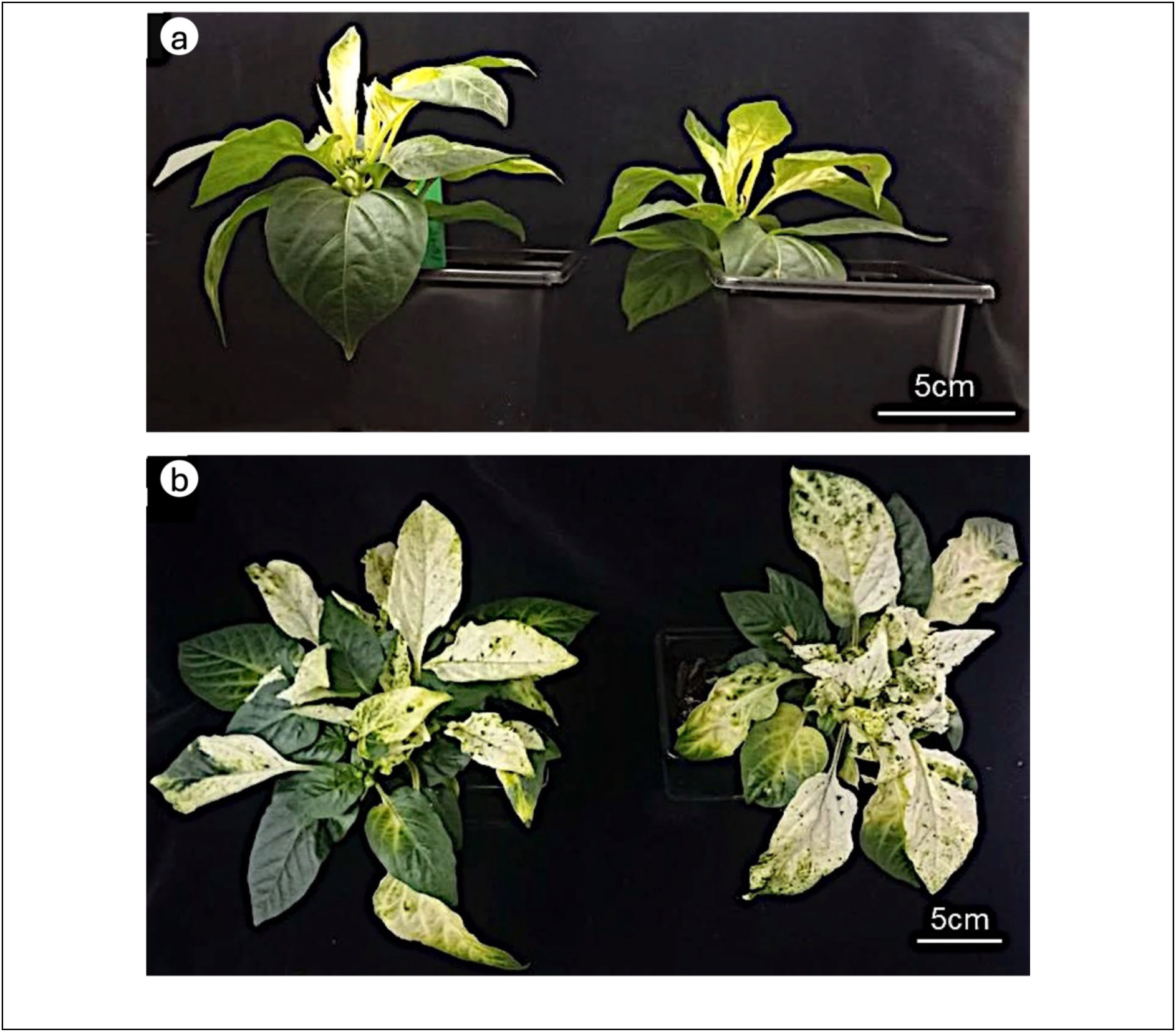
Agroinoculation of cotyledons was successful for launching TRV infection and induction of *PDS* silencing. Using Californian Wonder as an example, newly developing leaves on two plants show bleaching at six (a) and eight (b) weeks post-agroinoculation with a mixture of *Agrobacterium tumefaciens* cells carrying pYL192 (encoding TRV RNA1), pYL156:PDS (to silence *PDS*) and pYL156:BPEV (targeted against BPEV RNA).

**Supplementary Figure 6.**
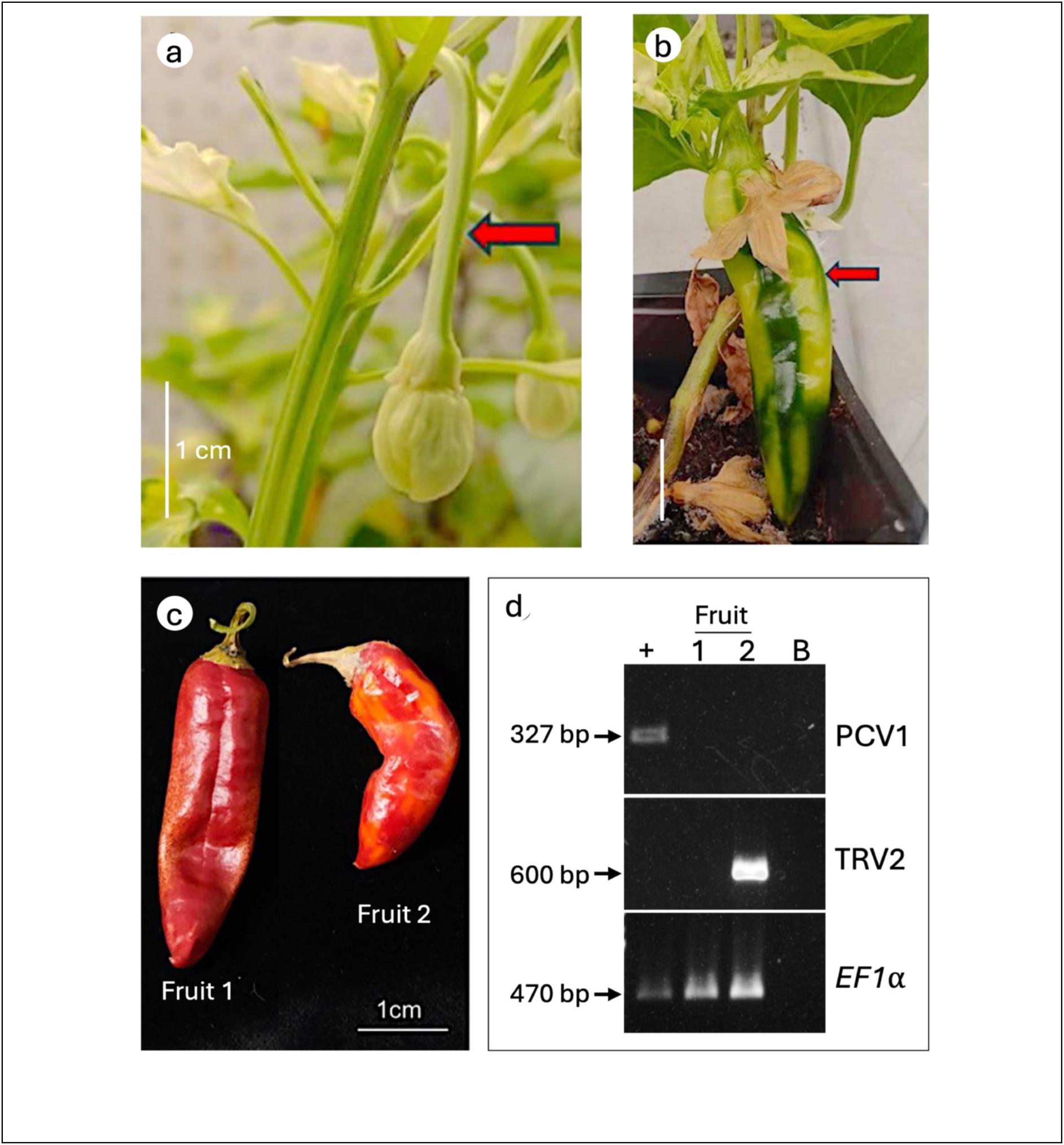
VIGS-mediated silencing of *PDS* and a persistent virus RNA in pepper. Using Mirasol Chilli as an example, pepper plants were agroinoculated in the cotyledons with TRV VIGS vector mixtures targeting *PDS* and PCV1 RNA1. *PDS* silencing resulting in photobleaching was apparent in flower peduncles (a) and fruit pericarp (b) (arrows). Photobleaching guided the selection of fruits and seeds for harvesting. Pericarp tissues from two ripened, bleached fruits (c) were also evaluated for PCV1 silencing (d). RNA extracted from bleached pericarps (Fruits 1 and 2) and a control fruit (+) from an untreated plant was used for RT-PCR with specific primers to test for PCV1 and TRV RNA2 (‘TRV2’). A control reaction was performed using primers for the pepper *EF1α* transcript. PCV1 was undetectable in fruit from plants subjected to *PDS* and PCV1 VIGS, while TRV RNA 2 was detected in one of the silenced fruits (Fruit 2) (c). Lane B was loaded with a sample of the no-template RT-PCR control reaction.

**Supplementary Figure 7.**
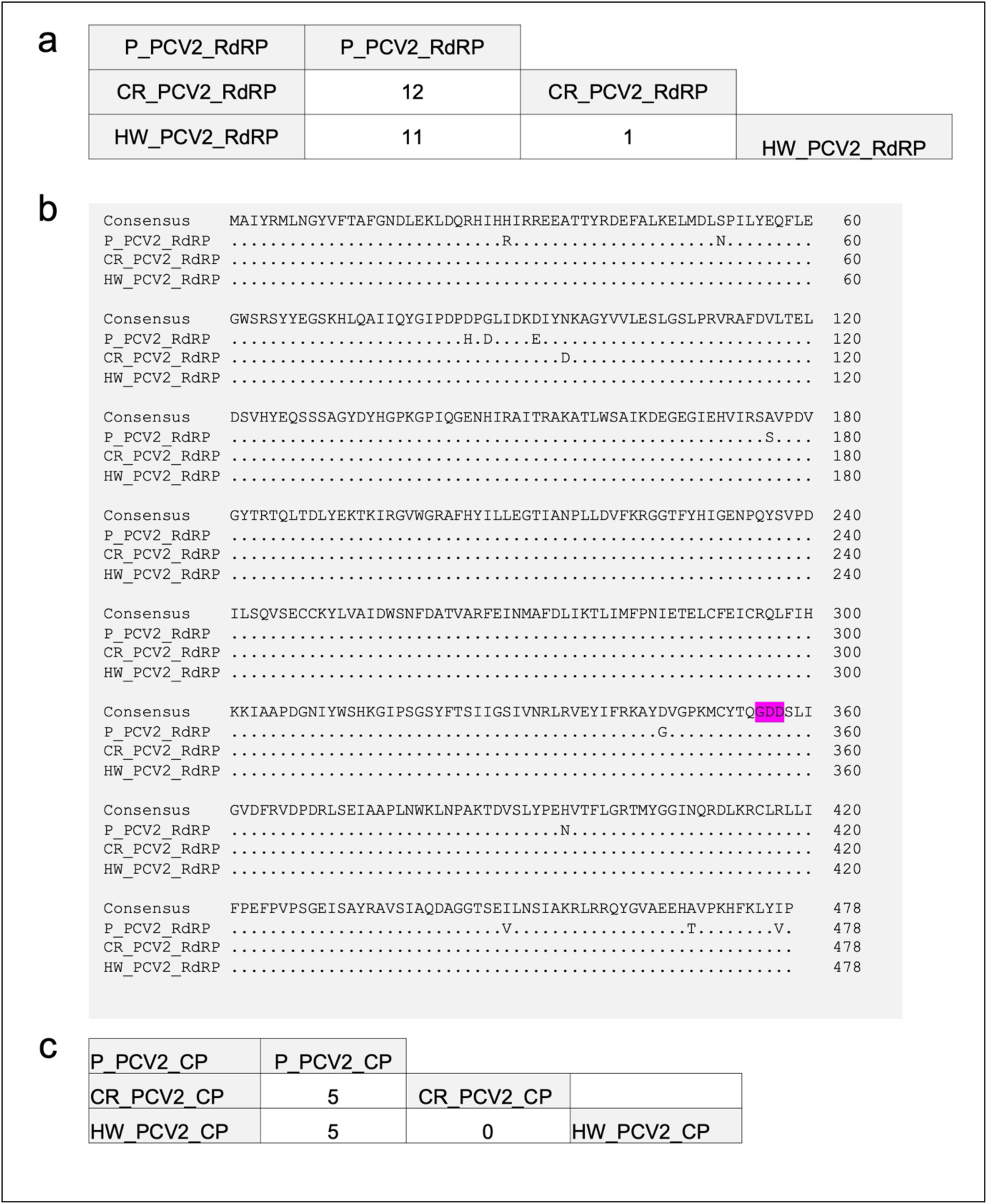
PCV2 variants from Hungarian Wax (HW) and Cayenne Red (CR) pepper cultivars are more similar to each other than to the PCV2 variant from Poupila **(P)** (a) Similarity matrix showing the number of amino acid differences between the RNA-dependent RNA polymerases (RdRP) encoded by the RNA1 open reading frames of the three PCV2 isolates. (b) Virtual translations of RdRP open reading frames from PCV2 RNAs 1 from Poupila, Cayenne Red, Hungarian Wax, with the conserved gly-asp-asp (GDD) box shaded. (c) Similarity matrix showing the number of amino acid differences between the coat proteins (CP) encoded by the RNAs 2 of the three PCV2 isolates.

**Supplementary Figure 8.**
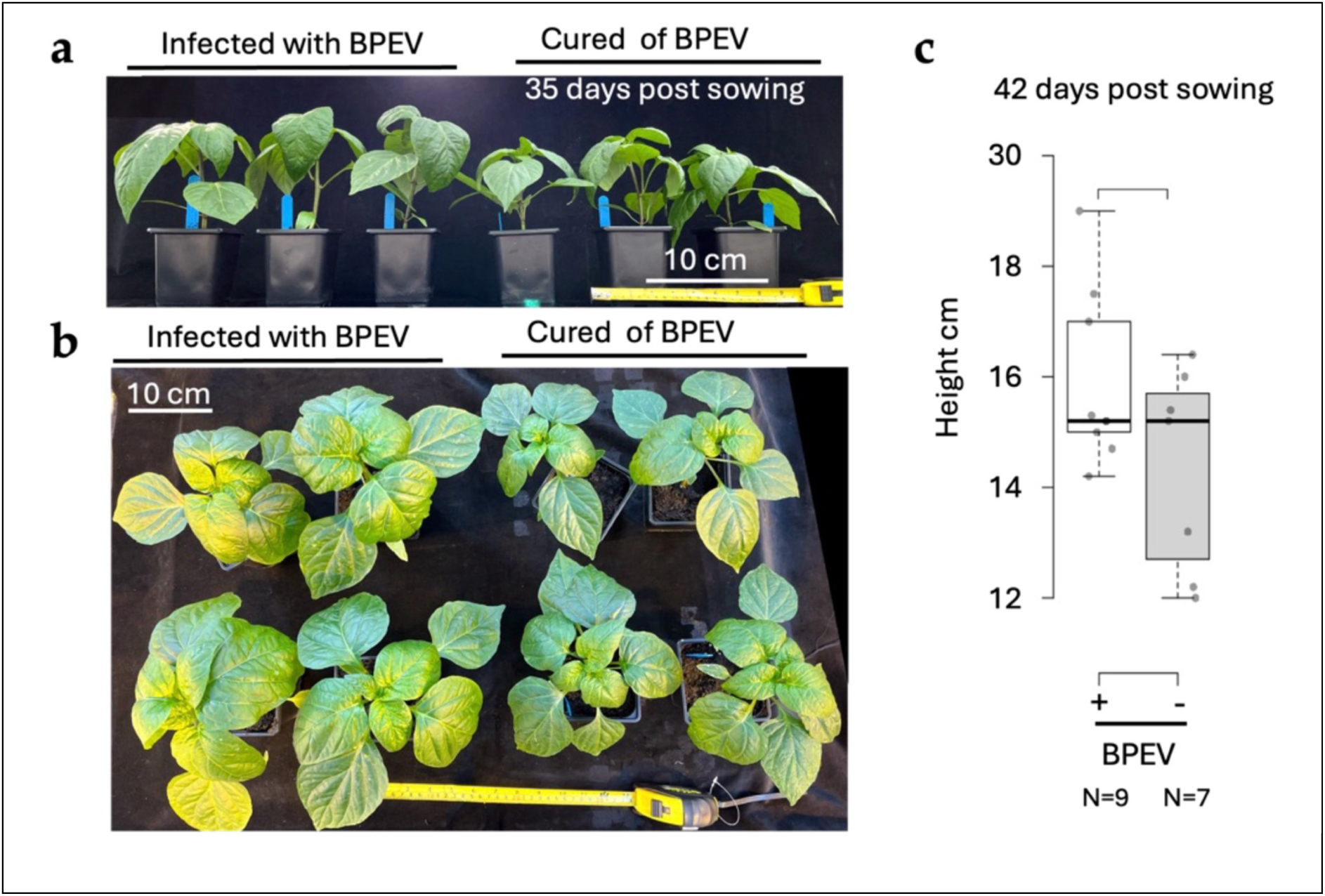
Growth of Californian Wonder harbouring and cured of BPEV. Seeds of plants silenced for BPEV (BPEV–) or of control plants infected with TRV-PDS (BPEV+) in the Californian Wonder background were germinated and the V1 plants (i.e., the progeny of plants subjected to VIGS) grown. Plant appearance was recorded 35 days post sowing (a, b) and plant height (measured to the apical bud) was recorded 42 days post sowing (c).

**Supplementary Figure 9.**
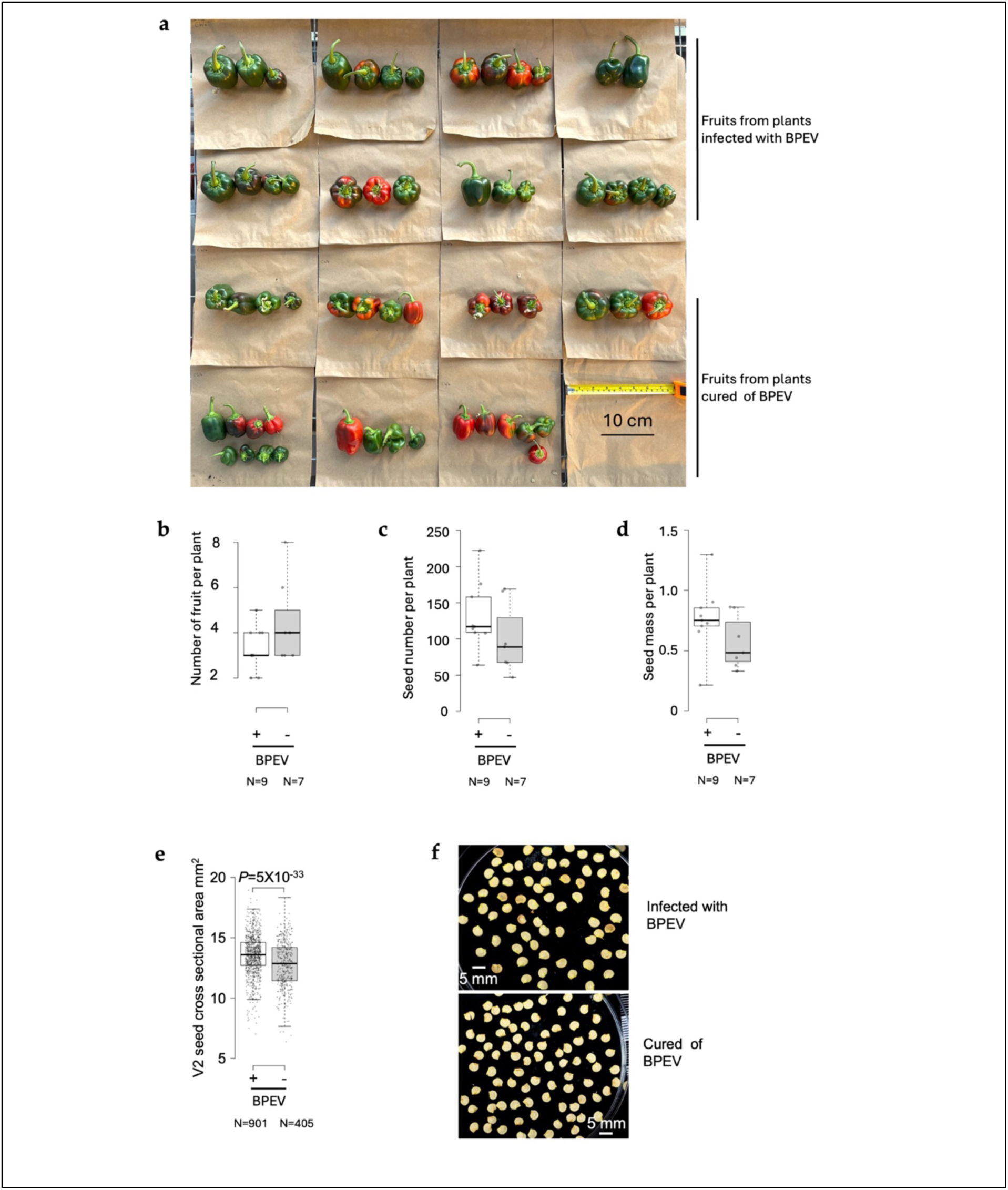
Fruit and seed production in Californian Wonder background plants harbouring or cured of bell pepper endornavirus (BPEV) Seeds produced from agroinfected plants either carrying BPEV (+) or cured of BPEV infection (–), were germinated and V1 plants (i.e., the progeny of plants subjected to VIGS) were grown to maturity and fruit harvested 127 days post sowing. No significant difference in fruit number was determined between BPEV-infected or cured plants (a, b). There were no significant differences in seed production per plant in terms of number (c) or mass (d) between BPEV-infected or cured plants. Mean seed size in this generation (‘V2’, i.e., the progeny of V1 plants and the second generation after VIGS) was significantly smaller in plants cured of BPEV compared to their BPEV-carrying counterparts (e, f).

**Supplementary Figure 10.**
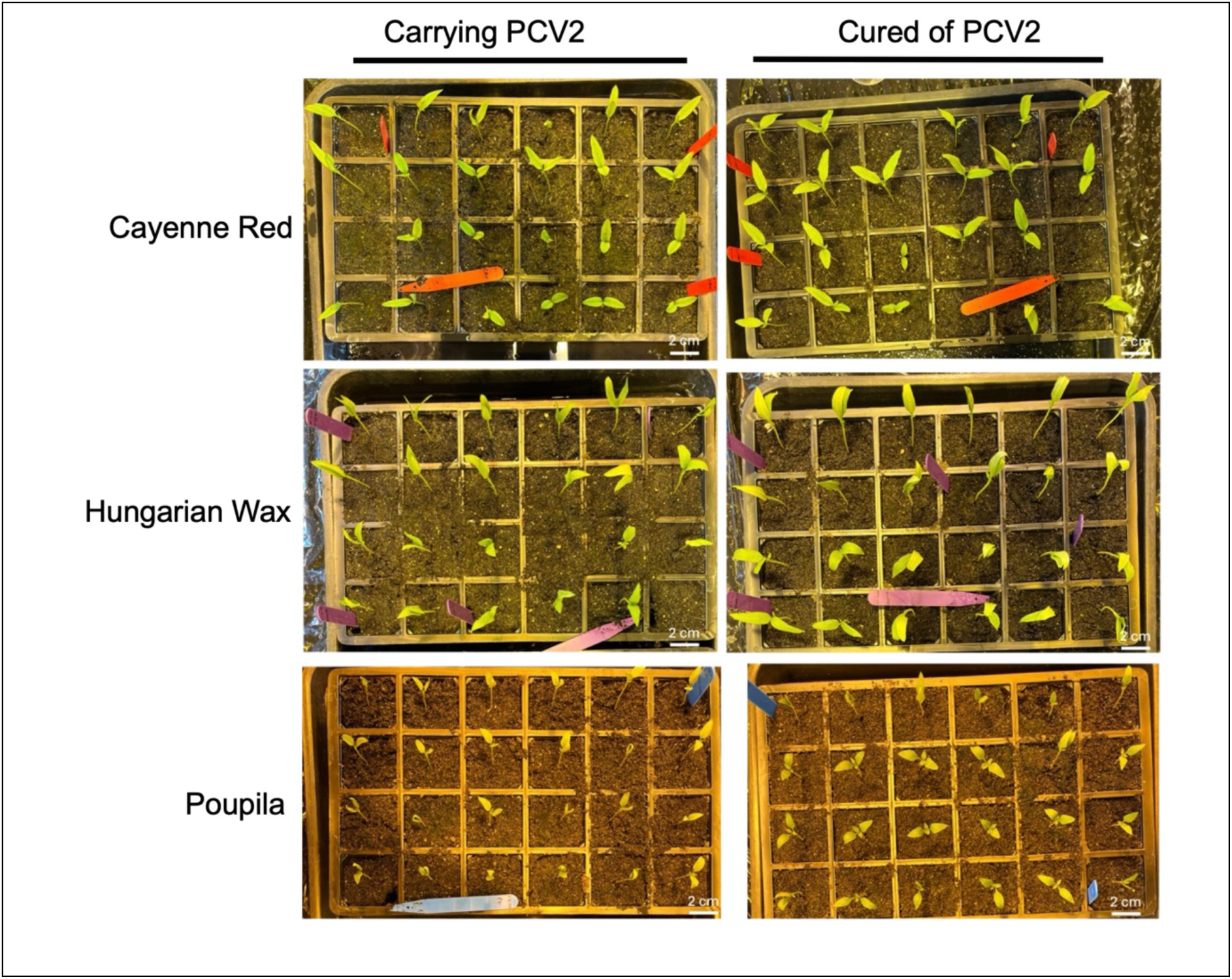
Establishment of peppers carrying or cured of PCV2. Appearance of V2 (i.e., the second generation after VIGS) seeds harvested from V1 plants (i.e., progeny of plants subjected to VIGS), either carrying or cured of PCV2 in three backgrounds (Cayenne Red, Hungarian Wax or Poupila), 14 days after sowing. Seeds germinated and successfully emerged at similar rates.

**Supplementary Figure 11.**
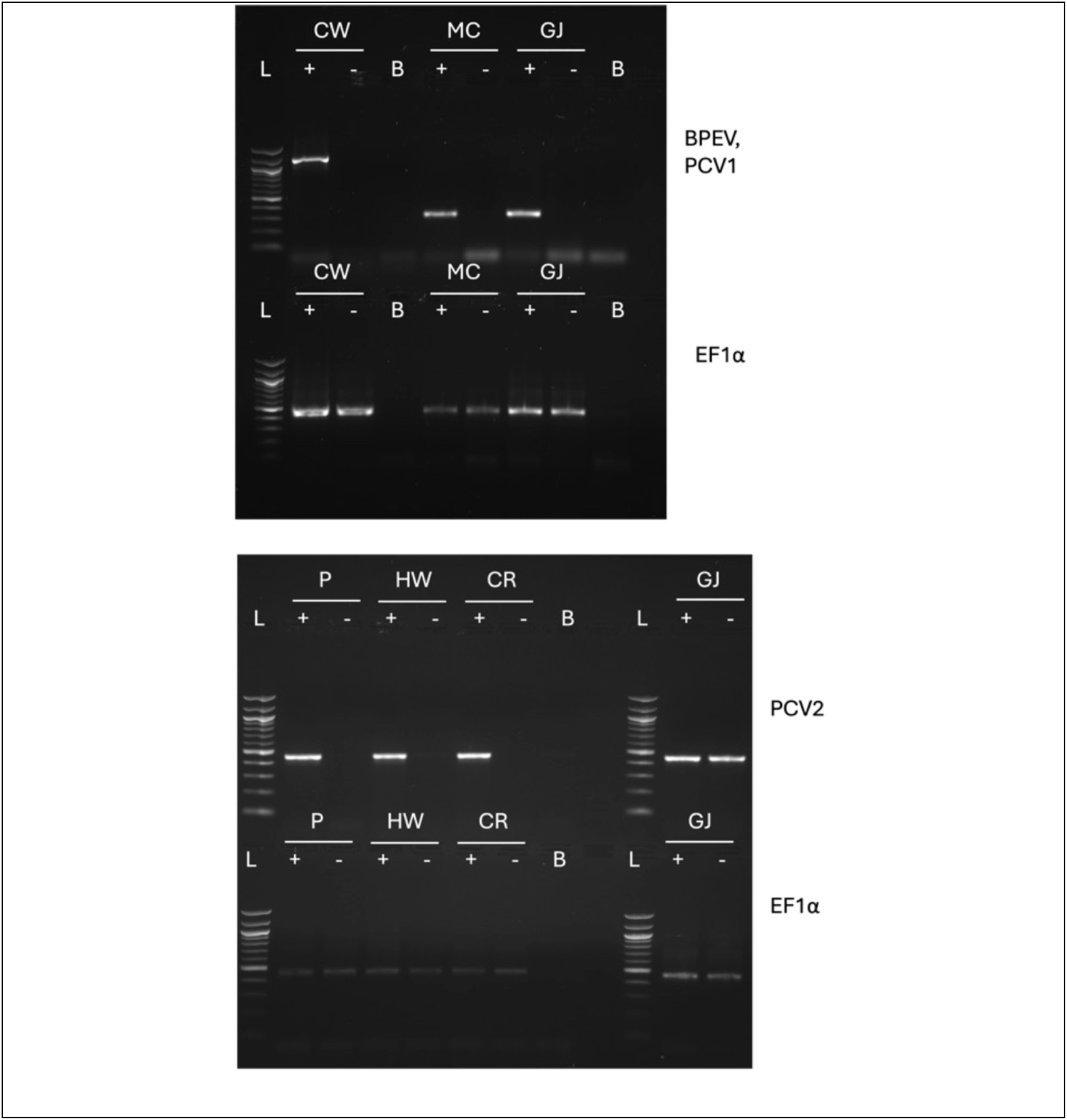
Original gel images used for Figure 2 (Silencing of persistent viruses in vegetative tissue of agroinfected pepper plants) RT-PCR using gene-specific primers targeting PCV1, PCV2, and BPEV was performed on leaf samples from agroinfiltrated pepper plants. Products were resolved on 1% agarose gels. Californian Wonder (CW), Mirasol Chilli (MC) and Gourmet Jalapeno (GJ) cultivars subjected to triple agroinfiltration show examples of silenced (-) and non-silenced (+) lines for BPEV and PCV1 (Top Panel). Poupila (P), Hungarian Wax (HW) and Cayenne Red (CR) subjected to triple agroinfiltration show examples of silenced and non-silenced lines for PCV2 (lower panel). Amplicons indicate non-silencing. In Gourmet Jalapeno infected with both PCV1 and PCV2, VIGS constructs containing pYL156:PCV1 successfully silenced PCV1 (upper panel) but left PCV2 unaffected (lower panel). Positive *CaEF1α* PCRs confirmed successful cDNA synthesis. Blank serves as a no-template control. L =100-bp DNA ladder.

**Supplementary Figure 12.**
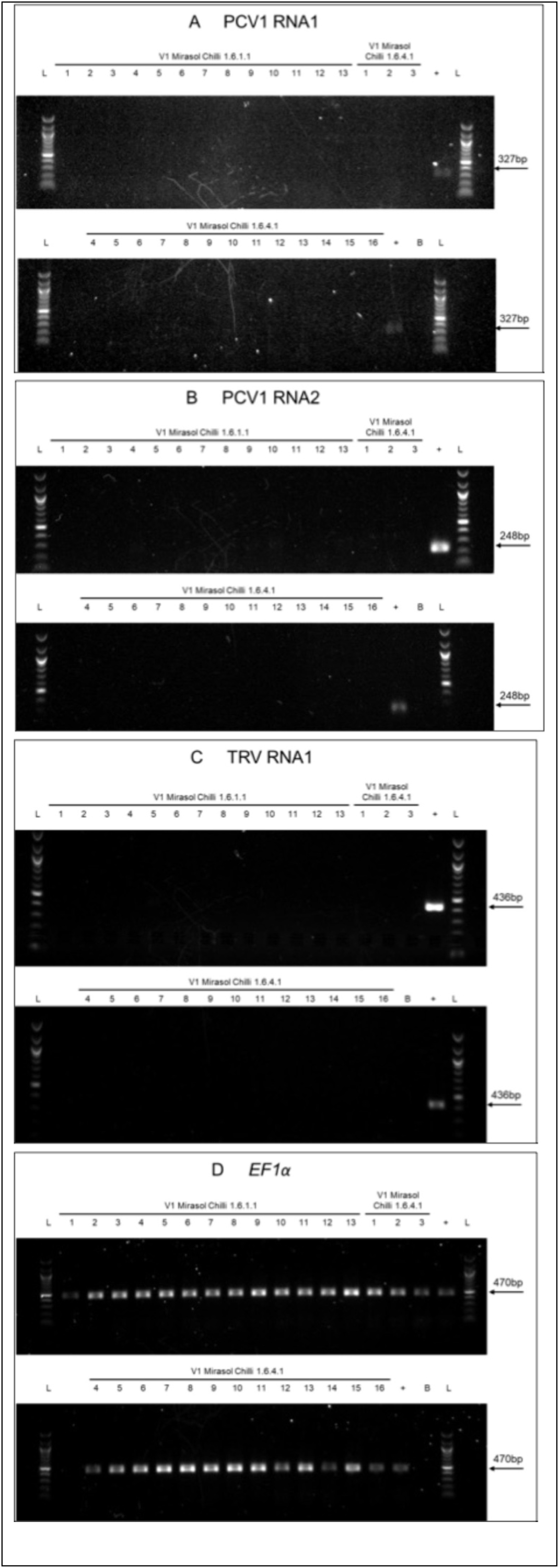
Original gel images used for Figure 3 (RT-PCR detection of viral RNAs in the progeny of silenced pepper plants) Reverse transcription followed by PCR with gene-specific primers targeting the PCV RNA1 and RNA2 and TRV RNA1 was carried out on tissue samples from pepper plants of V1 generation (i.e., progeny of plants subjected to VIGS) obtained from two PCV1 silenced Mirasol Chilli plants (1.6.1 and 1.6.4). RT-PCR products were separated on 1% agarose gels. TRV-PDS agroinfected pepper plants served as a positive control (+) for persistent virus and TRV infection. Numbers below the cultivars designate individual screened plants. Panels **A** and **B** show successful silencing of both RNA1 and RNA2 of PCV1 in the progeny of primary silenced pepper plants. Panel **C** shows that TRV is not inherited in the progeny of agroinfected plants. Panel **D** shows RT-PCR amplification of *CaEF1α,* which served as an internal control, confirming successful cDNA synthesis across all samples. The blank, no-template control (labelled B above lanes of gel) contained no cDNA in the PCR reaction mixture. A 100 bp DNA ladder (L) was included on each gel. Arrows indicate expected amplicon lengths.

### Supplementary Materials and Methods

#### Metatranscriptome library construction, high-throughput sequencing, non-host unigene assembly, and phylogenetic analysis

Total RNA was extracted from uncured plants belonging to the cultivars Californian Wonder (carrying BPEV), Mirasol Chilli (carrying PCV1), Hungarian Wax and Poupila (both carrying PCV2) and Gourmet Jalapeno (carrying PCV1 and PCV2) using Norgen Biotek Total RNA plus kits (Thorold, Ontario, Canada) that included removal of genomic DNA. Meta-transcriptome library preparation from the RNA samples and subsequent sequencing was carried out by the Novogene Company Limited, Cambridge, UK. Briefly, rRNA was depleted from total RNA to enrich mRNA and other non-ribosomal transcripts. The remaining RNA was reverse-transcribed to synthesize first-strand cDNA using random primers, followed by second-strand cDNA synthesis to generate double-stranded cDNA where dTTP was replaced by dUTP (for strand specificity). The cDNA fragments underwent end repair, 3′ adenylation, adapter ligation, size selection, and PCR enrichment to construct strand-specific libraries. Library quality and insert size distribution were assessed before sequencing on an Illumina NovaSeq X Plus Series platform using paired-end 150-bp reads. Sequencing was performed to a target depth of approximately 12 Gb of raw data per sample, generating paired-end reads of 150 bp from both ends of cDNA. Raw data underwent the Novogene workflow for quality filtering to remove low-quality reads and adapter sequences, retaining clean reads with a Q30 score > 85% for downstream taxonomic annotation. To isolate microbial (i.e. non-host) transcripts, reads were mapped against the *Capsicum annuum* reference genome assembly UCD10Xv1.1 (Hulse-Kemp et al., 2018; retrieved from the NCBI Genome Database under RefSeq accession GCF_002878395.1) using HISAT2 (hierarchical indexing for spliced alignment of transcripts 2; v2.0.5; Kim et al., 2019) to filter out host-derived sequences. The non-host reads were further filtered against the SILVA database using Bowtie2 to remove remaining prokaryotic and eukaryotic rRNA contamination. The remaining non-host reads were subjected to *de novo* transcriptome assembly using Trinity software (v2.15.2: Haas et al., 2013). To minimize sequencing noise and eliminate low-frequency base errors, the minimum *k*-mer coverage parameter was explicitly set to 2 (--min_kmer_cov 2), filtering out unique single-occurrence *k*-mers, while all other parameters were maintained at default settings. To eliminate sequence redundancy, the assembled transcripts were clustered using Corset (V1.09) to generate a set of non-redundant unigenes.

The non-host unigenes from each pepper cultivar were screened for viral sequences using BLASTn searches within Geneious Prime (version 2025.2.2) against a locally constructed NCBI viral genome database (retrieved from NCBI Virus, accessed May 2025). Non-host unigenes were also screened for viral hallmark sequence composition features using GeoNomad available at https://portal.nersc.gov/genomad/ (Camargo et al, 2024). No additional viral contigs were identified beyond those detected by prior BLAST-based screening, indicating that the persistent viruses identified in this study represent the complete set of detectable viral sequences, with no evidence of additional novel viruses.

Relevant full-length or near full-length reference genome sequences for PCV1, PCV2 and BPEV were retrieved from NCBI Virus (accessed May 2025; accession numbers listed in Supplementary Table 6) and were aligned with sequences from this study using Clustal Omega (with default settings) in Geneious Prime (version 2026.0.2, Biomatters Ltd). For each alignment, the best-fitting nucleotide substitution model for phylogenetic tree building was selected using the Model Selection tool in MEGA 12 (Kumar et al., 2024). Phylogenetic analysis was performed in [Geneious Prime](https://www.geneious.com/?utm_source=chatgpt.com) 2026.0.2 using the PhyML Maximum Likelihood algorithm where phylogenetic trees were inferred under the General Time Reversible model substitution model with gamma-distributed rate variation among sites (four rate categories) (Guindon et al., 2010). The gamma distribution parameter was estimated from the dataset during tree construction. Branch support was assessed using 1000 bootstrap replicates, and a consensus tree was generated. Branch lengths are proportional to the estimated number of substitutions per site. As the objective was to compare the relationships between newly sequenced persistent viruses with each other and related reference strains obtained from sequence databases, outgroup sequences were omitted. The final consensus trees were visualised and annotated within the Geneious interface.

